# Trinucleotide building blocks enable exponential ribozyme-catalysed RNA replication and open-ended growth of diverse RNA sequence pools

**DOI:** 10.1101/2023.03.17.533225

**Authors:** James Attwater, Teresa Augustin, Joseph F. Curran, Samantha Kwok, Edoardo Gianni, Philipp Holliger

## Abstract

RNA replication is considered a crucial stage in the origins of life. However, both enzymatic and non-enzymatic RNA replication cycles are impeded by the “strand separation problem” (SSP), a form of product inhibition arising from the extraordinary stability of RNA duplexes and their rapid kinetics of reannealing. Here we show that RNA trinucleotide triphosphates (triplets) can overcome the SSP by binding to and kinetically trapping dissociated RNA strands in a single-stranded form, while simultaneously serving as substrates for RNA replication by a triplet polymerase ribozyme (TPR). This enabled exponential replication of both (+) and (−) strands of double-stranded RNAs by the TPR when driven by coupled pH and freeze-thaw cycles. We demonstrate replication of a fragment of the ribozyme itself, and open-ended amplification of random RNA sequence pools over >70 cycles, with emergence of partial, distributive TPR self-replication and triplet codon drift towards a primordial genetic code.

**One-sentence summary:** RNA trinucleotide substrates together with simple physicochemical cycles enable RNA-catalysed replication of double-stranded RNA and partial, distributive self-replication of an RNA polymerase ribozyme.

## Main text

All life on Earth relies upon the faithful copying of its genetic material - replication - to enable heredity and evolution. This critical behaviour is widely thought to have begun with the templated polymerisation of activated mono- or oligo-nucleotide building blocks by chemical replication processes^1-3^ and later RNA-catalyzed RNA replication^4–7^ (Fig. 1a). In its simplest form RNA replication comprises the copying of (+) and (−) strands into complementary (−) and (+) daughter strands. For replication to proceed further, the double-stranded RNA replication products must be again dissociated into single-stranded RNAs and these daughter strands must be copied before they reanneal.

**Fig. 1.**
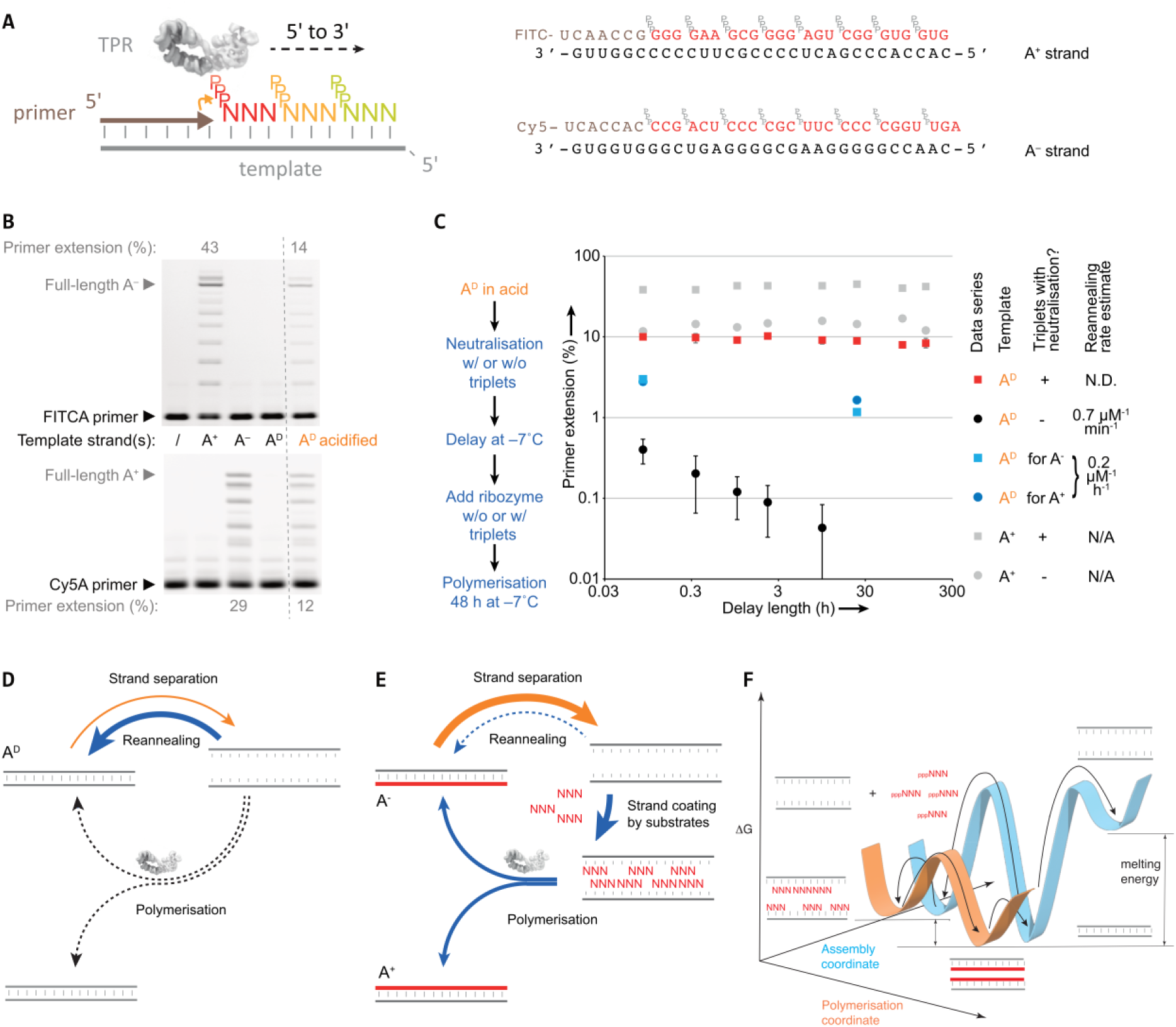
Triplet substrates alleviate strand reannealing during RNA replication. (**a**) Left, schematic of RNA strand copying by polymerisation of trinucleotide triphosphates (triplets) upon an RNA template, catalysed by a triplet polymerase ribozyme (TPR, structure from ref. 21). Right, substrates for synthesis of A^D^ duplex; individual strands (A^+^ and A^−^) are shown hybridized to their complementary primers (brown) and triplets (red). (**b**) PAGE gel of TPR RNA polymerisation with either A^D^ duplex or individual strands A^+^, A^−^ as templates showing synthesis of A^−^ (top, FITC channel)) and A^+^ (bottom, Cy5 channel). “A^D^ acidified” was pre-incubated in 2.5 mM HCl. After mixing in extension buffer (neutralizing any HCl), reactions were frozen (−7°C for 48 h). Shown are observed percentages of primer extended by >1 triplet in each reaction (after subtraction of levels in no-template controls (/)), measured by gel densitometry. (**c**) Effect on primer extension (%) of delaying triplet & primer addition after neutralisation of acidified A^D^ template. Reaction compositions (except template) were made identical prior to a polymerisation step in which primer extension was measured (black & red, n=3; error bars: ± s.d.). Curve fitting indicates that A^D^ reanneals with a t_1/2_ of 0.7 µM^-1^ min^-1^ (black circles). Addition of triplets upon A^D^ neutralization (red squares) essentially abolishes strand reannealing. (**d**) Challenges of RNA duplex replication. The difficulty of strand separation and ease of strand reannealing together inhibit RNA replication cycles. (**e**) Revised scheme of an RNA replication cycle driven towards polymerisation by triplet substrate inhibition of strand reannealing. (**f**) Schematic representation of the free energy landscape of triplet substrate assisted RNA replication.

However, attempts at recreating this fundamental process in the laboratory have been hindered by the challenge of having to overcome profound product inhibition by the duplex RNA products. RNA duplexes of useful lengths and concentrations (e.g. >25 nucleotides (nt), 100 nM) behave as “dead-end” products due to their remarkable stability (with melting temperatures that approach the boiling point of water)^8^. Furthermore, even when dissociated into individual strands, such RNAs re-anneal on timescales (seconds to minutes) that are considerably shorter than the typical timespan of copying reactions (hours to days) catalyzed either by non-enzymatic processes or by polymerase ribozymes^1^. Thus, RNA replication cycles under ambient conditions are both kinetically and thermodynamically disfavoured (Fig. 1d).

This so-called “strand separation problem”^1^ (SSP) is aggravated by the comparative chemical instability of RNA. This precludes dissociation under harsh conditions (such as high temperatures) that degrade RNA templates and ribozyme catalysts, particularly in the presence of divalent cations such as Mg^2+^ (which boost activity but accelerate the hydrolytic fragmentation of RNA oligomers)^9^. The SSP worsens with increasing lengths of RNA duplexes, which become progressively harder to dissociate, more vulnerable to degradation and more prone to reannealing.

Diverse approaches to overcome this fundamental barrier to RNA replication have been explored. For example, moderately acidic pH can protonate the N_1_ of adenine and N_3_ of cytosine bases, disrupting base pairing and destabilizing RNA duplexes^10^. Coupled with wet/dry cycles or ionic gradients in a thermophoretic setting, this has been shown to promote duplex melting and enable nucleic acid amplification by proteinaceous enzymes^11–14^. Furthermore, highly viscous solvents can slow RNA reannealing sufficiently for long (32-nt) substrates to be ligated^15, 16^. Circumventing full duplex dissociation, strand-displacement syntheses can occur on a short RNA duplex by the progressive addition of “invader” oligonucleotides complementary to the non-templating strand^17^, or by the buildup of conformational strain on circular RNA templates^18^. Despite these advances, the scope of RNA-catalysed RNA replication cycles has been limited: either via polymerisation of NTPs on primers flanking a 4-nt region (assisted by denaturants)^19^, or via templated ligation of (up to) three polynucleotide substrate segments^14, 20^. General RNA replication and open-ended evolution requires both the replication of longer sequences able to encode a phenotype and the polymerisation of building blocks short enough to enable free sequence variation.

Here, we describe a novel approach that leverages the properties of trinucleotide triphosphate (triplet) substrates together with cycles of opposing pH, temperature and concentration gradients to drive RNA replication by a triplet polymerase ribozyme (TPR). We show triplet-based exponential replication of both (+) and (−) strands of defined double-stranded RNAs, as well as > 70 cycles of open-ended amplification with serial dilution of random RNA sequence pools. Characterizing the replicated RNA pools by next-generation sequencing reveals both evidence of partial, distributive self-replication of the TPR itself as well as a sequence drift towards codon usage in an hypothetical primordial genetic code.

## Results

### Inhibition of strand reannealing by triplet substrates

We explored RNA replication using the 5TU / t1 polymerase ribozyme (a heterodimeric ribozyme^21^ henceforth referred to as ‘TPR’) that had been evolved *in vitro* to copy RNA templates using trinucleotide triphosphates (triplets) as substrates^4^ (Fig. 1a, Fig. S1). As shown previously, such template-directed RNA synthesis reactions by the TPR are preferentially carried out within the eutectic phase of water-ice at −7°C, due to high ionic and RNA substrate concentrations and reduced water activity^22^. Under these conditions triplet substrates display remarkable properties such as cooperative invasion of template RNA secondary structures, enabling copying of even highly structured RNA templates by the TPR^4^.

We initially hypothesized that this ability of triplets to invade and unravel template secondary structures might be leveraged to invade and replicate otherwise inert RNA duplexes. To test this we assembled a model 30 nt GC-rich RNA duplex A^D^ (Fig. 1a, predicted T_m_ = 99°C^23^) and incubated it together with constitutive triplet substrates and TPR. However, while the TPR readily synthesised full-length products on the duplex’s individual RNA (+) and (−) strands (A^+^ and A^−^), the A^D^ RNA duplex itself remained inert, as judged by the lack of primer extension activity (Fig. 1b). This indicates that - unlike RNA template secondary structures - triplets cannot productively invade an RNA duplex, at least under these conditions.

Next, we tested whether prior RNA duplex disruption might enable triplet invasion and replication by the TPR. Using a fluorescence-quench assay we found that temperatures >90°C were required to dissociate a mixed sequence 30 nt RNA duplex into constituent strands (Fig. S2). In the presence of mM concentrations of Mg^2+^ ions (needed for ribozyme catalysis) even higher temperatures (approaching the boiling point of water) were required. As exposure to such temperatures in the presence of Mg^2+^ would cause rapid hydrolysis and fragmentation of longer RNA strands, including the TPR itself, we explored alternative approaches to destabilize RNA duplexes. Mildly acidic pH had previously been shown to destabilize short RNA duplexes and is not destructive^10^ as RNA – unlike DNA – does not suffer depurination at moderately low pH. However, we found that at ambient temperatures duplex dissociation of >50% required a pH as low as 2.3 (with 20 mM MgCl_2_, or pH = 2.8 with 100 mM NaCl, Fig. S1). Though incompatible with polymerase ribozyme activity (pH_opt_ ≈ 8.8), we tested if acid-induced RNA duplex dissociation could be leveraged for copying of the A^D^ duplex as part of a sequence of: 1) acid denaturation of A^D^ duplex; 2) neutralisation and addition of TPR, primer and triplet substrates; and 3) freezing and polymerisation at −7°C. Indeed, this yielded full-length synthesis of both A^+^ and A^−^ constituent strands starting from A^D^ duplex template (at ∼40% of the yields compared to individual A^+^ or A^−^ strands as templates, Fig. 1b).

To better understand the observed simultaneous extension on the two complementary A^+^ and A^−^ strands, we investigated the kinetics of RNA duplex reannealing and replication in such reactions, by progressively delaying ribozyme addition after the neutralization step. To our surprise, extension on a pre-denatured duplex was maintained even when ribozyme was added up to 6 days post neutralization (Fig. 1c). However, if triplet addition was also delayed, subsequent extension was rapidly reduced (k_obs_ ∼0.7 µM^-1^ min^-1^) (Fig. 1c) presumably due to rapid reannealing of the two dissociated A^+^ and A^−^ RNA strands.

These observations are consistent with a scenario in which dissociated RNA strands become kinetically trapped in a single-stranded state when partially (or fully) hybridized to complementary triplets, which robustly attenuate strand reannealing, providing a time-window for RNA replication (Fig. 1e). Interestingly, this inhibition of intermolecular annealing is effective at lower triplet concentrations (∼2.5 µM/triplet, Fig. S3) than those needed for invasion of intramolecular template RNA secondary structures such as hairpins (>12 µM/triplet)^4^, suggesting the even partial triplet occupancy may be sufficient to attenuate re-annealing. Indeed, even addition of triplets complementary to just one of the strands (A^+^ or A^−^) slowed re-annealing by ∼200-fold (Fig. 1c), while addition of complementary triplets to both strands effectively stops the second-order re-annealing process and maintains strands in a dissociated state for >300 h. Thus, triplets appear to play a dual role, both as RNA chaperones keeping complementary strands from re-annealing, and as substrates for RNA replication on the dissociated strands. Such “substrate-assisted replication” is wholly contingent on the nature of the triplet substrates, as an RNA polymerase ribozyme using mononucleotide triphosphate (NTP) substrates^24^ exhibited negligible extension on the same denatured duplex (Fig. S4).

Mechanistically, it seems likely that triplets exert their effect by 1) base-pairing to and selectively stabilising transient “open” states of RNA, 2) progressively “coating” RNA strands via specific hybridization and cooperative triplet-triplet base stacking interactions, and 3) acting as substrates for TPR-catalyzed synthesis of a complementary RNA strand. While 1) is an equilibrium process, 2) and 3) are effectively irreversible, ratchet-like processes, which produce progressively longer RNA products further stabilizing the template dissociated state until a full complementary strand is synthesized. Furthermore, we observed that the TPR has the capacity to initiate templated RNA synthesis from adjacent triplets at multiple sites along the template (Fig. S5). Such “primer-free” RNA replication both supports the above mechanistic model and likely accelerates 3) complementary strand synthesis. Therefore, by blocking strand reannealing and creating a long-lived substrate:template complex, triplets serve to decouple RNA polymerisation parameters from both the kinetics and thermodynamics of RNA duplex dissociation and reannealing (Fig. 1e, f).

### Iterative RNA replication cycles

Having discovered an effective strategy to overcome the strand separation problem, we sought to integrate triplet-assisted strand separation and triplet-based RNA synthesis into a full RNA replication cycle. However, the preferred conditions required for strand separation and RNA replication are diametrically opposed. Optimal conditions for RNA duplex denaturation include low pH and elevated temperatures to weaken base-pairing interactions together with low Mg^2+^ concentrations ([Mg^2+^]) to minimize RNA hydrolysis and duplex stability, and low RNA strand concentrations to attenuate reannealing upon neutralization (Fig. S6). In contrast, optimal RNA replication conditions include a mildly basic pH, high [Mg^2+^], high ribozyme and triplet substrate ([RNA]) concentrations and ambient to low temperatures. Furthermore, incubation of RNA at acidic pH in the presence of the high [Mg^2+^] needed for optimal ribozyme activity (> 60mM) can cause precipitation and inactivation of long RNAs - an effect exploited in the trichloroacetic acid (TCA) precipitation of nucleic acids. A replication cycle would therefore require opposite, reversible shifts in pH, temperature and solute concentrations ([Mg^2+^], [RNA]) (Fig. 2a).

**Fig. 2.**
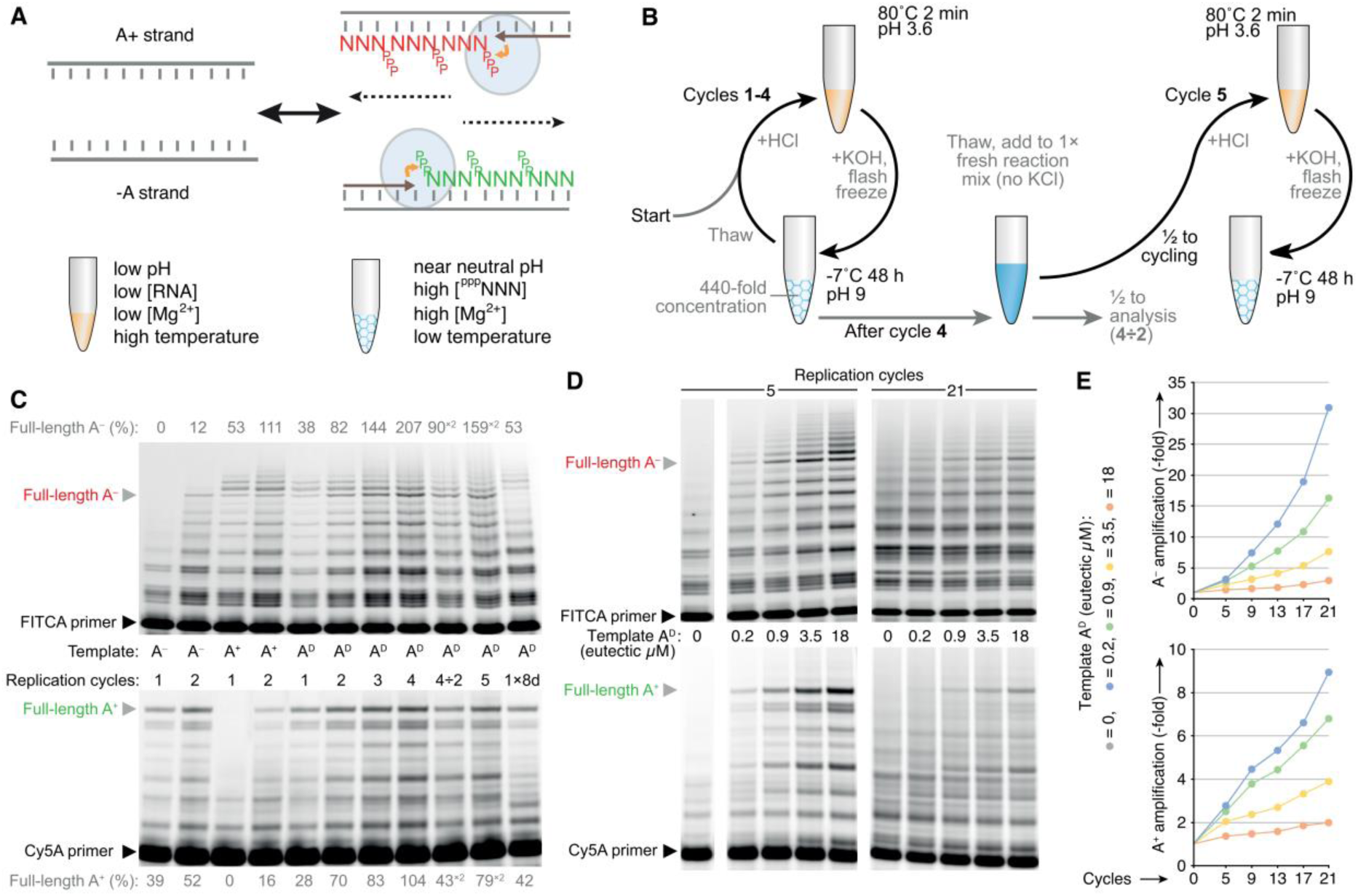
Freeze/thaw pH cycling allows iterative RNA replication. (**a**) Schematic of conflicting conditions required for RNA strand separation (left) and triplet polymerisation (right). (**b**) Physicochemical cycling workflow integrating strand separation and polymerisation conditions. pH switching results in a build-up of KCl; serial dilution allows continued cycling by resetting KCl concentrations and restoring ribozyme and triplet substrate levels. (**c**) Iterative replication of the model RNA duplex A^D^ and its constituent strands. Denaturing PAGE of extension products after cycles in replication buffer (0.4 mM MgCl_2_, 1 mM CHES·KOH pH 9.0, 0.01% Tween-20) with 1.8 mM KCl, 20 nM TPR, 4 nM template, and 0.1 µM of each substrate and primer as in Fig. 1a. Also shown are four-cycle reactions undergoing two-fold dilution (4÷2) followed by an extra cycle (5), and (for comparison) a single cycle 8 day polymerization reaction (1×8d). Full-length primer extension efficiencies in each lane are measured by densitometry and expressed as percentages relative to starting template. (**d**) Denaturing PAGE of A^D^ replication reactions set up as in (c) with varying starting A^D^ duplex concentrations, continued over 21 cycles including 5 serial dilutions. (**e**) Calculated product amplification in reactions in (d). The amount of primer reaching full-length (above that in the no template control) was observed by densitometry and normalized to the levels of A^D^ after serial dilution.

To begin to reconcile these conflicting requirements, we first defined milder, more dilute denaturing conditions (pH 3.6, 80°C, low [Mg^2+^] & [RNA]) that efficiently separate even GC-rich RNA duplexes (Fig. S2) whilst avoiding RNA degradation. To move to synthesis conditions, lowering temperature and increasing pH are easily achieved; the key challenge is to drive concentration shifts to high [Mg^2+^] & [RNA]. We had previously discovered that eutectic ice phases were an ideal environment for polymerase ribozyme activity, and that freezing can drive a dramatic concentration of solutes and reduction of water activity within the eutectic brine phase^22^. Therefore, after optimizing replication buffer composition together with neutralization and freezing conditions to maximize free strand availability (Fig. S6), we defined a coupled pH / freeze-thaw cycling protocol (Fig. 2b), which was capable of coupling denaturing conditions that efficiently dissociate RNA duplexes (Fig. S1, S6) with extension conditions that yield near-full TPR activity (Fig. S7).

Next we tested whether this strategy of coupled pH / freeze-thaw cycles could support multiple iterative rounds of replication and yield amplification of double-stranded RNA sequences. In a single cycle starting from individual strands, A^−^ or A^+^ templated full-length RNA primer extension by the TPR using defined substrates with per-strand yields of 39% or 53%, and a second cycle up to doubled these yields (52% or 111%), as measured by gel densitometry (Fig. 2c). Uniquely, this second cycle was also accompanied by full-length extension of primers corresponding to the starting strands (yields of 12% (A^−^) and 16% (A^+^)). This indicates that in cycle 2, the product strands of cycle 1 are used as templates, thereby providing a foundation for exponential RNA amplification. Next, we initiated repeated cycles of pH / freeze-thaw replication starting from the duplex RNA A^D^ and observed simultaneous production of both full-length A^−^ and A^+^ synthesis products. After four cycles product yields reached approximately two A^−^ strands (207%) and one A^+^ strand (104%) per starting A^D^ duplex (compared to ∼0.5 strand of each in a single long cycle with equivalent total polymerisation time) (Fig. 2c).

### Open-ended RNA amplification

In our pH / freeze-thaw cycling scheme, HCl and KOH addition drives the pH changes, but results in a build-up of K^+^ (and Cl^-^) ions, eventually inhibiting polymerisation (Fig. S6). We solved this by imposing a serial, two-fold dilution regime (into fresh reaction mix containing ribozyme and substrates, but lacking template and KCl) every fourth cycle to reset ionic strength (Fig. 2b). Continuing A^D^ replication under this regime, 21 cycles yielded diminishing returns in A^+^ and A^−^ yields (Fig. 2d), presumably due to RNA degradation and the low synthetic yield of A^+^ strands. Nevertheless, we observe effective strand amplifications of 9-fold (A^+^) and 31-fold (A^−^) after accounting for the overall 32-fold reaction dilution from the lowest starting levels of the template duplex A^D^ (Fig. 2e).

To better assess the potential of this cycling protocol in an open-ended context, we next sought to apply it to a diverse pool of RNA sequences using random triplet substrates (all 64 ^ppp^NNN). We initiated RNA replication from an RNA sequence pool comprising an 18 nt random-sequence region (N_18_) flanked by defined primer binding sites. We hypothesized that, if the RNA replication protocol was functional, we would observe drift and eventually persistence of those RNAs via amplification that can both be synthesized and act as efficient templates for replication (Fig. 3a).

**Fig. 3.**
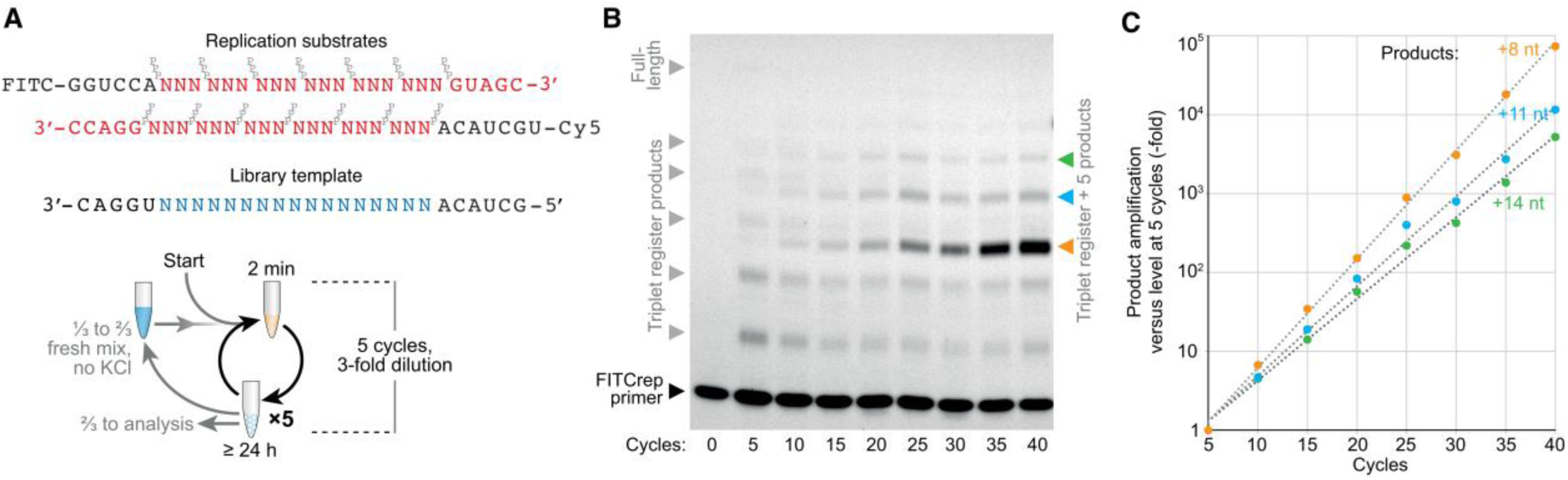
Open-ended RNA amplification. (**a**) Design of replication substrates and scheme for iterative replication of an N_18_ RNA random-sequence library. To start, library template LibN_18_ was mixed at 8 nM in replication buffer (including 0.9 mM KCl and 20 nM TPR) together with 20 nM each of the indicated primers (FITCrep, Cy5rep, ^ppp^GUAGC, ^ppp^GGACC) and 12 nM each of all 64 triplets (^ppp^NNN). (**b**) Denaturing PAGE of FITCrep extension products from this reaction analysed at 5-cycle intervals before 3-fold serial dilution. (**c**) Quantification of overall amplification of ‘triplet register +5’-length products in part B, calculated as fold increase in band intensity since 5 cycles, multiplied by reaction dilution since 5 cycles. Exponential fits yielded observed per-cycle amplification efficiencies.

In early cycles, a ladder of triplet register extension products was generated corresponding to the expected library size (6 × ^ppp^NNN triplet incorporations + a terminal pentamer (^ppp^GUAGC) adding a primer binding site, Fig. 3b). As cycling progressed (up to 40 cycles), this full-length product progressively faded, and a second series of extension products emerged and increased up to 30-fold in abundance, despite a >2000-fold effective dilution (an overall 6 × 10^4^-fold amplification). These +8, +11 and +14 nt product classes were identifiable by their altered register of migration and exponential amplification (1.37-, 1.3- and 1.27-fold per cycle respectively, Fig. 3c). These truncated product classes likely reflect early incorporation of the library’s terminal pentanucleotide substrate, providing a reciprocal primer binding site, and supporting higher yield synthesis of the shorter amplicons – a persistent phenomenon in PCR-style primed amplifications^25^. Nevertheless, the exponentially increasing yield of these emergent product classes confirms that with pH / freeze-thaw cycling the TPR can amplify a pool of RNA sequences in an open-ended one-pot reaction.

### Fragmentary self-replication

We next wondered if replication could be extended to the TPR’s own sequence. We had previously shown that the TPR can synthesize segments of both its (+) and (−) strand and re-assemble the (+) strand segments into a catalytically active ribozyme^4^. We tested the 29 nt *γ* fragment of the ribozyme catalytic core (Fig. 4a), confirming that without cycling both *γ*^+^ and *γ*^−^ strands can be synthesized by the TPR (Fig. 4b). A major concern in this context of self-replication was the risk of mutual inhibition of replication by the complementarity of TPR ribozyme and template. Specifically, we were concerned that the *γ*^−^ strand templates would invade the TPR structure during denaturation and become bound to the TPR *γ*^+^ region. However, one cycle of replication on *γ*^D^ efficiently yielded both full-length *γ*^+^ and *γ*^−^ product strands (Fig. 4b), and after two cycles we saw evidence of exponential replication on both individual *γ*^+^ / *γ*^−^ and duplex *γ*^D^ templates (Fig. 4b, c) with products being used again as templates in a second cycle as observed for A^D^ (Fig. 2c). This suggests that during one-pot denaturing cycles the TPR tertiary structure is either resistant to denaturation or refolds with faster kinetics than intermolecular invasion by complementary strands can occur. After five cycles of *γ*^D^ replication, per-duplex yields of extension products reaching full length exceeded 220%, consistent with a 1.26-fold amplification per cycle (Fig. 4d).

**Fig. 4.**
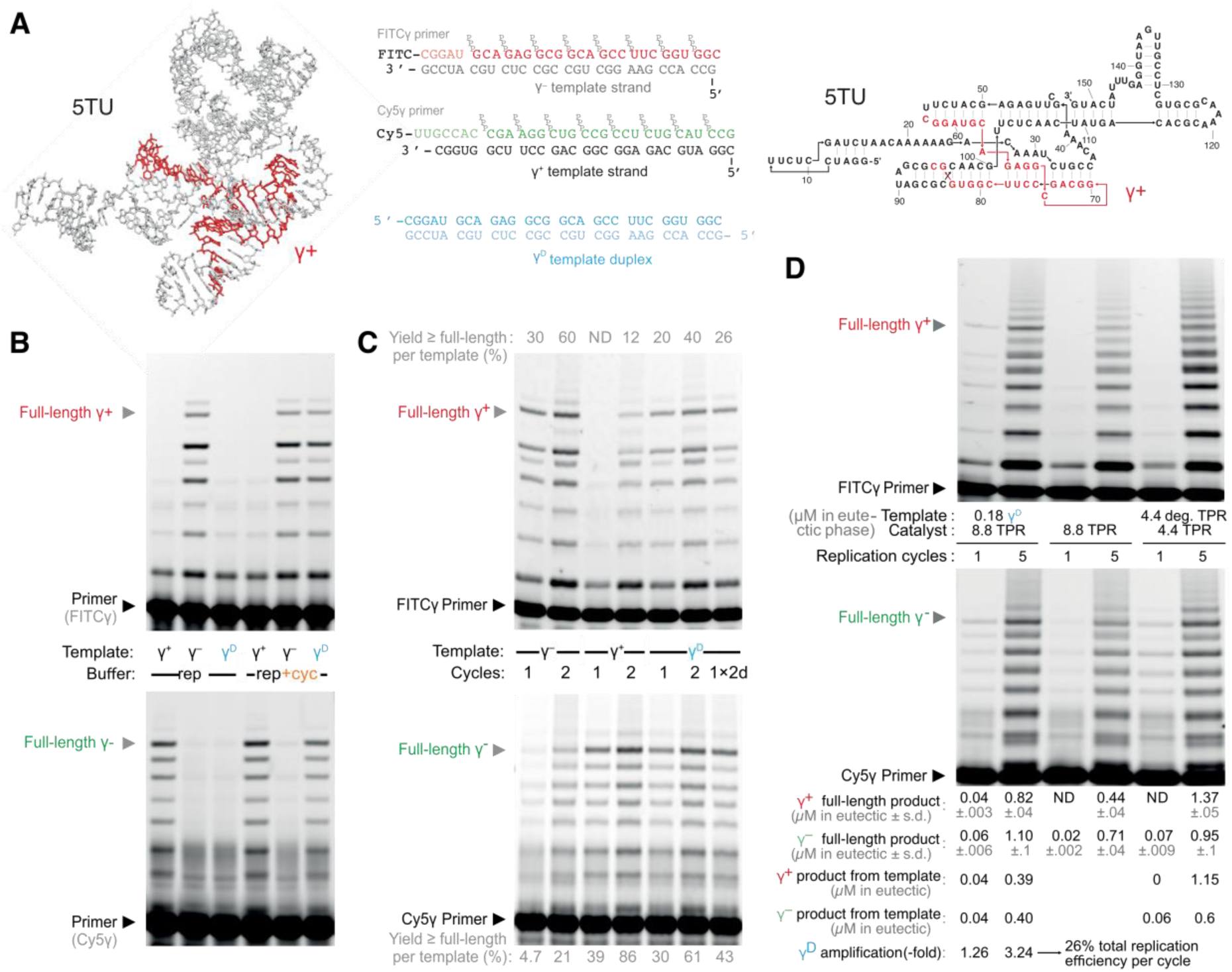
Ribozyme-catalysed replication of a fragment of itself. (**a**) 3D structural model^21^ (left) and secondary structure (right) of the 5TU catalytic subunit with the γ^+^ segment highlighted (red). Middle, sequences of the γ^D^ duplex and its constituent strands γ^+^ and γ^−^, shown with primer and triplet substrates. (**b**) Denaturing PAGE of γ^+^ and γ^−^ synthesis from γ^D^ template. The triplets (0.2 µM each) and primers (0.1 µM each) shown in (a) were polymerized (24 h) by the TPR (20 nM) upon the indicated templates (2 nM each) in replication buffer (‘rep’, with 2.4 mM KCl), with and without a preceding denaturing acid cycle.. (**c**) Primer extension (as in (**b**)), for one or two replication cycles. Per-template yields of primer reaching full-length (determined via gel densitometry) are given, showing reciprocal γ^+^ strand copying after two cycles, and TPR-templated γ^-^ strand synthesis after one cycle (at ∼1% the efficiency of templating by γ^+^). Two cycles yielded more product than a single cycle with equivalent total incubation time in ice (1×2d). (**d**) Iterative self-replication of the γ fragment by the TPR. Replication reactions were set up as in (**b**) (with 0.6 mM KCl), and either seeded with γ^D^, degraded TPR (by prior heating with Mg^2+^ = ‘deg TPR’), or no template. Post-freezing (eutectic phase) concentrations of template, TPR and products are given. Concentrations of γ^+^ (red) and γ^−^ (green) products reaching full-length were inferred from gel densitometry (n=3). Concentrations of product derived from replication of the template seed were calculated by proportional subtraction of products derived from intact ribozyme in unseeded reactions.

Unexpectedly, we also observed a different, asymmetric interaction of the TPR with the replication reaction, whereby unseeded, no template negative controls also yielded *γ*^+^ and *γ*^-^ product strands. Even after a single cycle, both unseeded or exclusively *γ*^−^-seeded reactions yielded *γ*^-^ products (whilst *γ*^+^-seeded reactions showed no *γ*^+^ synthesis) (Fig. 4c). We hypothesized that the *γ*^+^ segment within the TPR itself might be acting as a template (although at low efficiency (∼1%) compared to exogenously added *γ*^+^ template, Fig. 4c). Indeed, when we intentionally partially degraded 50% of the TPR (by heating at pH 9.0 with Mg^2+^) prior to replication cycling, we observed *γ*^+^ and *γ*^-^ synthesis products with yields comparable to *γ*^D^-seeded reactions, and specifically *γ*^-^ products after a single cycle (Fig. 4d). This suggested that the TPR fragments produced by partial degradation can act as templates, boosting product yields despite the lower amount of TPR catalyst available to drive RNA replication.

Sequencing the products of *γ*^D^-seeded reactions confirmed the accurate replication of γ^+^ sequences, alongside a number of partially-homologous sequences likely derived from incomplete extension products, and even some unanticipated products complementary to parts of the TPR t1 subunit (Fig. S8). Weighting the sequence data by reaction yield gave estimates of generation rates of different sequence classes, establishing that the generation rate of accurate γ^+^ products increased almost 50% from cycles 1-5 of γ^D^-seeded replication, with a 130% replication yield of γ^+^ strands after five cycles (Fig. S8). Lower levels of accurate γ^+^ sequences also emerged in the unseeded reaction. Our results thus show that 1) a ribozyme can exponentially replicate part of itself from short building blocks, 2) this ‘fragmentary’ self-replication can occur in a ‘one-pot’ cycled reaction and can 3) initiate even in the absence of a seed template using the TPR ribozyme itself.

### Emergence and replication of RNA sequence pools

In the replication reactions described so far (Fig. 2-4), we provided sequence-specific RNA primers to either target replication of defined sequences or random sequences of a defined length. However, the availability of sequence-defined RNA oligomers able to act as primers cannot be assumed *a priori* in a primordial context^1^. The capacity of the TPR to initiate replication from triplets internal to a template (Fig. S5) could remove the need for primers, potentially providing a more realistic primer-independent model of RNA-catalyzed RNA replication in which the system is free to explore RNA sequence space in an unsupervised manner.

To explore this behaviour, we initiated open-ended RNA replication by the TPR, without primers but with all 64 triplet substrates (^ppp^NNN) and an N_20_ random-sequence RNA seed (Fig. 5a). After five cycles a ladder of triplet-register RNA products was detected (Fig. 5b), indicating primer-free RNA synthesis from the N_20_ template seed. Surprisingly, a similar (if fainter) pattern was also seen in an unseeded control, implying emergence of products even in the absence of a seed template. Upon continued cycling (with serial dilution), levels of synthesised RNA in both seeded and unseeded reaction pools grew substantially and product lengths increased: after 73 cycles (and over 2.5 × 10^5^-fold dilution of the initial reaction), a robust ladder of products up to 36 nt in length was present in both seeded and unseeded reactions (Fig. 5b), consuming a substantial fraction of the ^ppp^NNN triplet substrates supplied upon each dilution step (Fig. 5c). While the seeded reaction initially maintained a higher level of RNA products (over a 128-fold dilution), RNA product levels in the unseeded reaction eventually caught up to the seeded reaction.

**Fig. 5.**
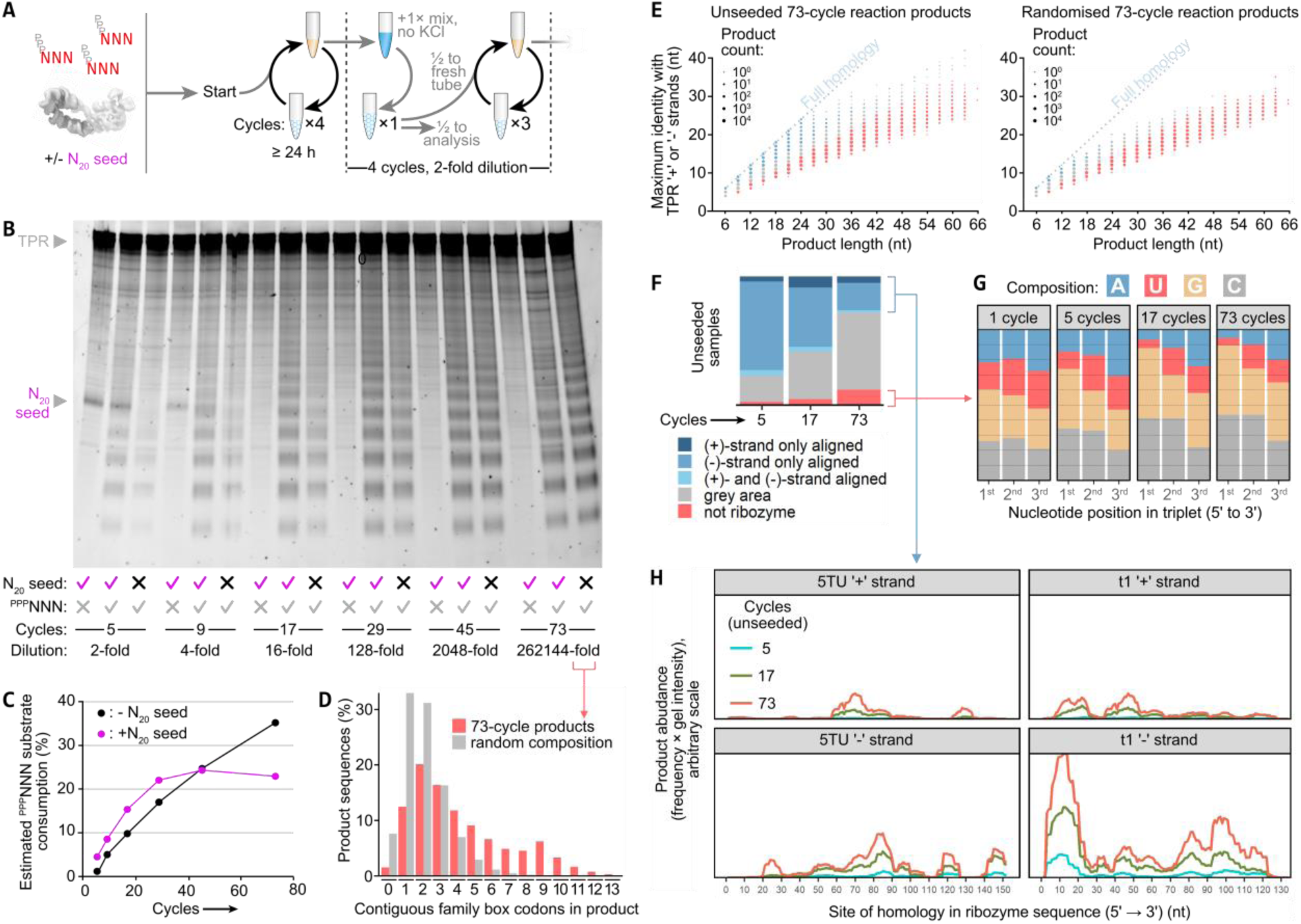
Emergence and maintenance of RNA pools upon primer-free triplet-based replication. (**a**) Replication reactions were set up with or without 0.1 µM of all 64 RNA triplets (^ppp^NNN) in replication buffer (no primers, 20 nM TPR, 1.8 mM KCl) ± 20 nM N_20_ random-sequence RNA template seed, and subjected to iterative cycles of replication and dilution as shown. (**b**) Denaturing PAGE of RNA samples at different stages (up to 73 cycles) stained with SYBRgold: both intensity and length of RNA products increased during cycling and serial dilution. (**c**) Estimated conversion of the total triplet substrate pool into RNA products. Lane intensities in (b) beneath the TPR bands were measured by densitometry, and the corresponding ^ppp^NNN-free reaction backgrounds (and, if present, N_20_ seed band intensities) were subtracted. These intensities were converted to RNA product yields using the N_20_ seed band intensity as a reference ((b), left lane), treating fluorescence of stained RNAs as proportional to their length. The triplets needed to make these RNA yields were expressed as a percentage of the available triplet substrate. (**d**) In silico ‘translation’ using a reduced codon set of sequenced unseeded 73-cycle synthesis products (red) compared to a simulated pool of random sequences with matching lengths but unbiased composition (grey). For each sequence the longest stretch of family box codons is counted to show the maximum potential length of any encoded peptide at a primitive stage of genetic code development. (**e**) Ribozyme sequence complementarity in sequenced synthesis products from the unseeded 73-cycle reaction (left) and a simulated pool of randomised RNAs of identical composition (right). Data coloured by classification in (f) (see Fig. S9 for criteria used); sequences with homology to ribozyme (+) strand are plotted separately (Fig. S9). (**f**) Changes in proportions of sequence classes in 9-27 nt products from unseeded reactions during amplification. (**g**) As cycling progresses, the G-C base composition of sequences classed as having no ribozyme homology increases (data from N_20_-seeded reactions shown). (**h**) Mapping of ribozyme-homologous parts of 9-27 nt products from unseeded amplification reactions to the (+) and (−) strands of the TPR subunits 5TU and t1. Peak heights reflect the fraction of products homologous to that site, scaled by the product intensity in the corresponding gel lane in (**b**). Products with homology to multiple locations on one or both strands were randomly assigned to one. Note the prior emergence and build up of (−) strand TPR homology products, followed by (+) strand products (templated from (−) strand products).

To understand the nature of the RNA sequences that emerged in these reactions, we performed high throughput sequencing of all 5’-triphosphorylated RNAs from both seeded and unseeded reactions. In both, a substantial fraction of products (25 - 75%) showed evidence of partial complementarity to the ribozyme sequence (Fig. 5e, f, Fig. S9), suggesting that – as in the primer-dependent γ^D^ replication (Fig. 4d) – the TPR acted both as a polymerase and as a template in their generation. The homologous regions could be mapped to multiple points along the TPR sequence (Fig. 5h), presumably initiated from ligation of adjacent triplets as observed in model primer-free synthesis experiments (Fig. S5). Though some TPR sequence segments appear to be absent (e.g. 5TU (−) 0-20, 130-140), amplification products with sequence segments complementary to a TPR subunit (5TU / t1 (−) strands) build up progressively in the cycling reactions. We also observed a lower level of products with homology to 5TU / t1 (+) strands. Though some of these might arise through background sequencing of ribozyme catalyst, the majority had only partial homology to TPR suggesting that the emergent (−) strand products begin to act as templates themselves instructing (+) strand synthesis (Fig. 5h, Fig. S9). Therefore, randomly triplet-primed RNA replication driven by pH-freeze-thaw cycles may offer a spontaneous path to distributive TPR self-replication.

A significant fraction (5-20%) of RNA sequences generated in both N_20_-seeded and unseeded reactions showed no evidence of homology to either TPR (+) or (−) strands, and therefore are likely to have arisen by a different process. These sequences were extremely diverse (∼95% unique) (Fig. S10) and lacked demonstrable complementarity among themselves (Fig. S10). Nonetheless, across iterative replication cycles their abundance increased, and compositionally shifted towards a more GC-rich pattern (Fig. 5g) reminiscent of the TPR’s substrate preferences^4^. However, unlike a previous template selection experiment (using a mononucleotide RNA polymerase ribozyme), where products became progressively biased towards G-rich sequences^24^, here the relative proportions of G to C as well as of A to U remained both stable and closely matched, analogous to Chargaff’s rule^26^. Thus these emerging RNA sequences are likely propagated in a templated process^27^. Their emergence in both seeded and unseeded reactions suggest that in addition to random self-priming and distributive replication, the TPR spontaneously generates a pool of novel RNA sequences which – in contrast to products of random sequence amplification using primers (see Fig. 3) – grow longer and more complex upon open-ended cycling. Thus, ribozyme-catalysed replication with random-sequence substrate pools would be accompanied by both emergent distributive self-replication as well as a “creative mode” driving *de novo* generation of diverse RNA sequence pools.

## Discussion

Here, we have explored the capacity of RNA trinucleotide triphosphates (triplets) to provide an efficient solution for the strand separation problem of RNA replication. By binding and stabilizing RNA oligomers in single-stranded form, triplets can attenuate strand reannealing. At the same time, template-bound triplets can serve as both substrates and initiators (primers) of RNA replication by a triplet polymerase ribozyme (TPR). Leveraging these properties in combination with coupled pH, temperature and concentration gradients enabled replication and exponential replication of defined double stranded RNA sequences and random RNA sequence pools by the TPR. Furthermore, the ability of triplets to randomly initiate RNA synthesis enabled primer-free amplification of random RNA sequence pools resulting in de novo sequence generation as well as partial, distributive self-replication, with the TPR acting as both catalyst and template.

Cyclic variations in pH and temperature and ice formation shown here to drive RNA replication can be found in present day geothermal fields^29^ and are plausibly present on the early earth^28^. Furthermore, functionally analogous physicochemical gradients may be provided by a range of divergent geochemical environments including evaporation-condensation cycles or thermophoretic pH and concentration gradients^13, 30, 31^. Freeze-thaw cycles specifically had previously been shown to promote RNA folding^32^, and in recent reports was also shown to enable both activation and non-enzymatic RNA polymerisation^2^ as well as RNA 2’,3’-aminoacylation^33^. As the processes of strand separation and stabilization described here are driven by the physicochemical properties of RNA oligomers in a non-equilibrium setting and not by the nature of the catalyst, the same principles for overcoming the strand separation problem likely apply in non-enzymatic RNA replication^1^.

Despite the capacity of RNA triplets to hybridize to RNA strands and prime RNA synthesis as well as invade RNA secondary structure^4^, we find that the TPR remains surprisingly active in the presence of random triplet (^ppp^NNN) pools even during denaturing cycles (Fig. 5), presumably due to either the stability or rapid refolding kinetics of the TPR tertiary structure. Indeed, the TPR appears largely resistant to duplex sequestration with a complementary internal segment (γ^−^), allowing ‘one-pot’ self-replication of this fragment of itself (Fig. 4). However, the γ segment is part of the catalytic core of the TPR and it is currently not clear if less stably folded, more flexible segments of the TPR (such as the 5TU 3’ domain) would be more vulnerable to sequestration. Indeed, one might speculate that this property might be inversely related to the ability of different parts of the TPR to serve as templates. A future RNA replicase ribozyme will therefore have to manage the tradeoffs that come with embodying both template and catalysts in one.

One way to manage such tradeoffs is already evident in our replication experiments, where the TPR molecules initially serve as only inefficient templates, though with increasing efficiency as the TPR begins to unfold and/or degrade upon prolonged cycling (Fig. 4d). As replication cycles proceed, triplet priming on such fragments creates a growing pool of first TPR (−) and then (+) strand-homologous products, providing a potential route to an emergent form of distributive self-replication^34^ (Fig. 5h). Beyond these TPR fragment pools, replication with all 64 triplet substrates also results in the *de novo* generation and amplification of diverse pools of RNA oligomer sequences (up to ∼30 nt long). Elucidating their origins will require further study, though a variety of generational mechanisms have been identified in an analogous context catalysed by a proteinaceous polymerase^35^. Such sequences could serve as raw material for evolution - for example, by potentiating TPR activity or founding new activities to promote their own survival.

Analysis of the amplified product pools also reveals a striking drift (∼80%) towards sequences composed of triplets corresponding to family box codons in the genetic code (Supplementary Tables 3 & 4). These (1/2 of all codons) encode amino acids independently of their 3^rd^ nucleobase. Such triplets generally form more stable codon-anticodon interactions^36^, and together encode a putative amino acid set available from prebiotic chemistry^37, 38^ and are therefore widely thought to have been part of an simpler, primordial genetic code. Their selective enrichment in TPR replication products may reflect both TPR sequence preferences (for GC-rich triplets), as well as thermodynamic considerations (i.e. higher template occupancy) that have been shown to influence sequence pool evolution in model replication experiments^11^. In this way, triplet-based RNA replication embodies physicochemical principles that may also have shaped the early genetic code. Thus, by causing a drift in triplet usage, replication cycles shape sequence composition of random RNA sequence pools towards productive translation by a proto-ribosome utilizing family box codons. Indeed, the reduced incidence of unused codons would favour production of a population of longer peptides (Fig. 5d), potentially aiding the emergence of coded peptide based phenotypes from such sequence pools.

In summary, our results suggests mechanisms by which a nascent primordial biology could reconcile the conflicting roles of RNA as a general substrate, a replication template and a folded, stable catalyst. Our work defines physicochemical parameters needed for overcoming the strand separation problem and suggests that in the right environment, partial self-replication – at the level of sequence fragments in a distributive replication model^27, 34^ – appears as a predisposed, emergent property of RNA replication. Coupling this to a selectable phenotype may enable the maintenance and ultimately evolution of function in a future fully-synthetic replication system.

## Supplementary Figures

**Fig. S1.**
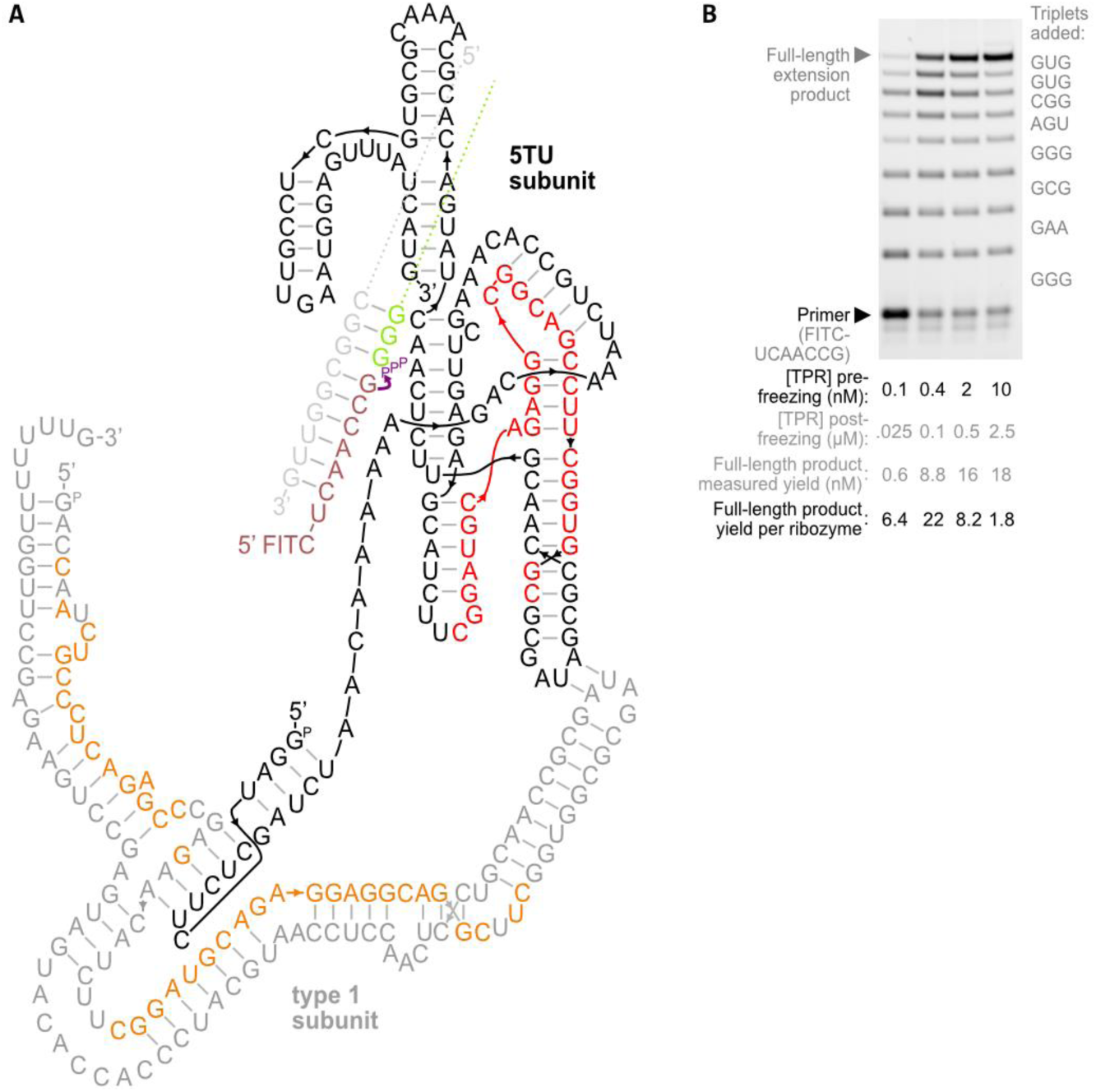
Features and activity of the triplet polymerase ribozyme 5TU^+1^ used in this study. (**a**) A secondary-structure-level representation of the 5TU^+1^ triplet polymerase ribozyme heterodimer based upon its cryo-EM structural model^21^. The TPR comprises a catalytic type 5 subunit (black) and an inactive type 1 subunit (grey), depicted surrounding a template-substrate complex. Triplet substrate (green) binding to template (light grey) juxtaposes the 3’ hydroxyl of the upstream RNA primer (brown) and the 5’ triphosphate of the triplet (purple). The type 5 subunit catalyses nucleophilic attack of the 3’ hydroxyl on the triphosphate α-phosphate of a correctly-paired triplet to form a new phosphodiester linkage. Type 1 forms a heterodimer with type 5, improving type 5’s interaction with ligation junctions. The γ fragment of 5TU is highlighted in red, and type 1 regions complementary to products in Fig. S8 are highlighted in orange. (**b**) Primer extension by the TPR. Full-length product is generated when all junctions are ligated, and incomplete ligation yields a ladder of intermediate extension products. Here, catalysis by the TPR synthesises complementary strand RNA upon multiple template copies. 40 nM of template A^+^ and primer FITCA were mixed with the indicated concentrations of TPR in 1 mM MgCl_2_, 1 mM CHES·KOH pH 9.0, 4 mM KCl, 0.001% Tween-20, and 0.1 µM of each triplet substrate. To initiate extension, the mixture was frozen in dry ice then incubated at −7°C for 24 h. Here, solutes are concentrated by an estimated 250-fold within the resulting supercooled liquid brine eutectic phase. Reactions were subsequently thawed, ethanol precipitated, resuspended in 6 M urea and heated to separate labelled products from template prior to resolution by denaturing PAGE. The percentage of fully-extended primers was then measured by gel densitometry, allowing calculation of number of complementary strands synthesised per ribozyme molecule (with each strand itself the product of eight iterative ligation reactions).

**Fig. S2.**
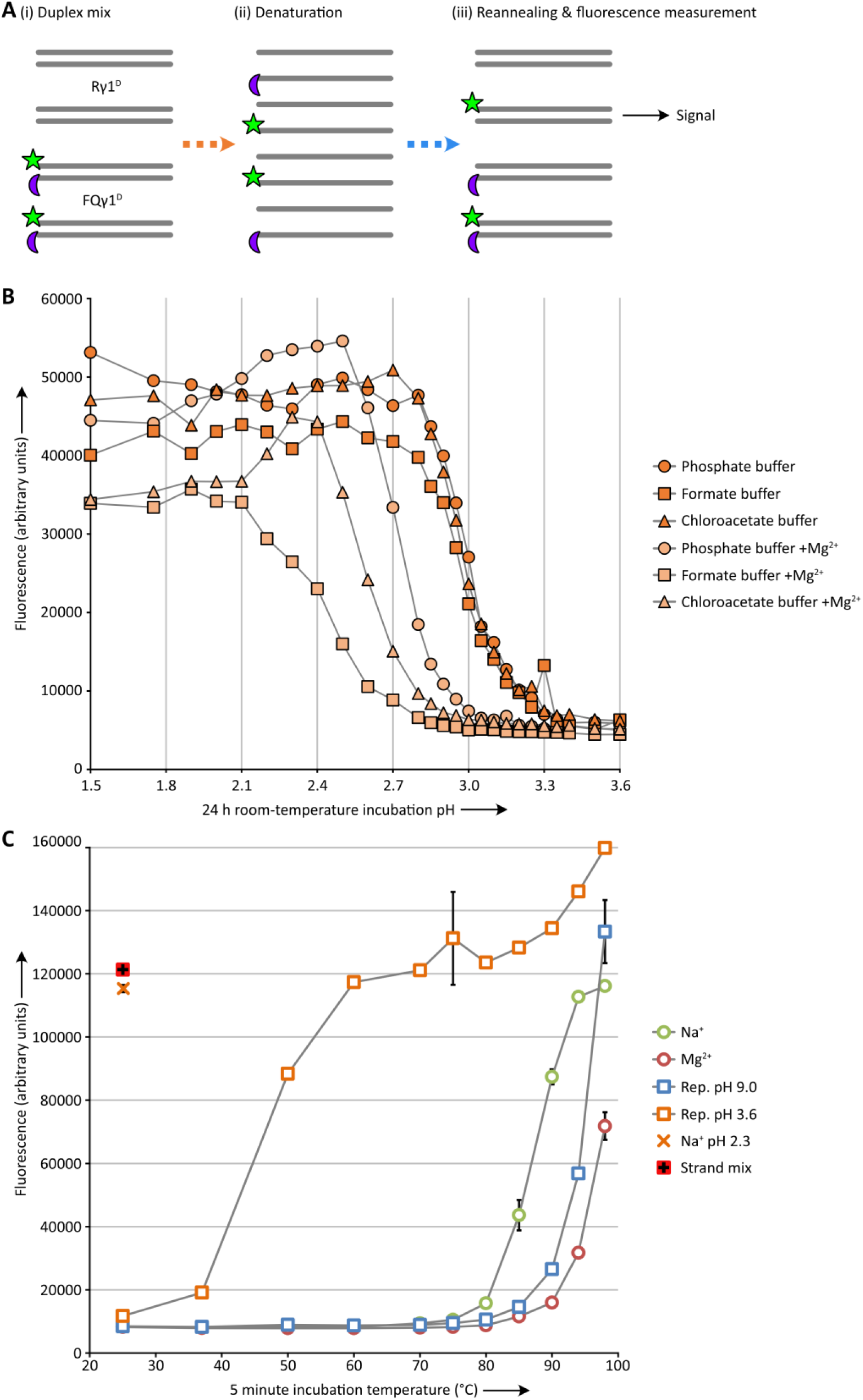
Fluorescence/quench assay of strand separation conditions. (**a**) Assay design. (i) The 27-nt RNA duplex Rγ1^D^ (70% GC composition) is mixed with an equal amount of duplex FQγ1^D^, whose component strands are identical but for a 5’ fluorescein group (green) on one and a 3’ Black Hole Quencher group (purple) on the other. Each duplex had been purified by native PAGE before mixing, and FQγ1^D^ yields low fluorescence due to the adjacency of the fluorophore and quencher on the hybridized strands. (ii) Upon subjecting the duplex mix to denaturing conditions, the strands separate from their starting partners. (iii) Returning to native conditions (pH 8, room temperature) leads to reannealing and reassortment of strands. Some fluorophore-labelled strands are now hybridised to unlabelled complements, yielding fluorescent signal in proportion to the extent of duplex dissociation. This assay decouples the conditions required for measurement from those needed for strand separation, and thus measures the history of denaturation in a sample, not its current state. (**b**) Strand separation of GC-rich RNA duplexes at low pHs. 0.5 pmol each of Rγ1^D^ and FQγ1^D^ were mixed in 40 µl of 0.1 M NaCl, 0.05% Tween-20 detergent and 0.1 M of phosphate, formate or chloroacetate buffers at the indicated pHs. After 24 h at room temperature, half of each sample was mixed with 1.5 volumes of 1 M tris·HCl pH 8, and its fluorescence measured using a Pherastar plate reader. Raw fluorescence (ex. 488 nm/em. 525 nm) of each sample is plotted against incubation buffer pH, demonstrating that lowering the pH below 3 can drive a sharp strand separation transition at room temperature. This is substantially below the pK_a_s of the nucleosides adenosine (4.5) and cytidine (4.0), potentially reflecting pK_a_ shifts when Watson-Crick base paired, and the need for cooperative protonation to disrupt a long duplex. A second set of samples (+Mg^2+^) with 20 mM MgCl_2_ also present in the buffer during denaturation showed a requirement for even lower pHs to separate the strands; furthermore, we observed that 20 mM MgCl_2_ at such pHs led to aggregation of longer RNAs such as the triplet polymerase ribozyme. (**c**) Strand separation of GC-rich RNA duplexes by heating. 1 pmol each of Rγ1^D^ and FQγ1^D^ were mixed in 50 µl of 0.02% Tween-20 and the following buffer compositions: 0.1 M NaCl, 1 mM tris·HCl pH 7.4 (Na^+^); 0.1 M NaCl, 20 mM MgCl_2_, 1 mM tris·HCl pH 7.4 (Mg^2+^); 2.4 mM KCl, 0.4 mM MgCl_2_, 1 mM CHES·KOH pH 9.0 (Rep. pH 9.0); 2.4 mM KCl, 0.4 mM MgCl_2_, 0.25 mM HCl (Rep. pH 3.6); 0.1 M NaCl, 5 mM HCl (Na^+^ pH 2.3). The samples were incubated for five minutes at the indicated temperatures, then cooled on the bench and neutralized with 50 µl 1 M tris·HCl pH 8, 25 mM EDTA. After sitting overnight at room temperature, sample fluorescence was measured in triplicate (error bars represent ± s.d.s) and plotted against incubation temperature. A positive control (Strand mix) comprising a mix of 1 pmol of both (+) with 1 pmol of both (−) strands of the duplexes in ‘Na+’ buffer conditions was also prepared. Very high temperatures are needed to dissociate duplex under neutral or slightly basic conditions, but at an acidic pH of 3.6 moderate heating is sufficient to denature the RNA duplex.

**Fig. S3.**
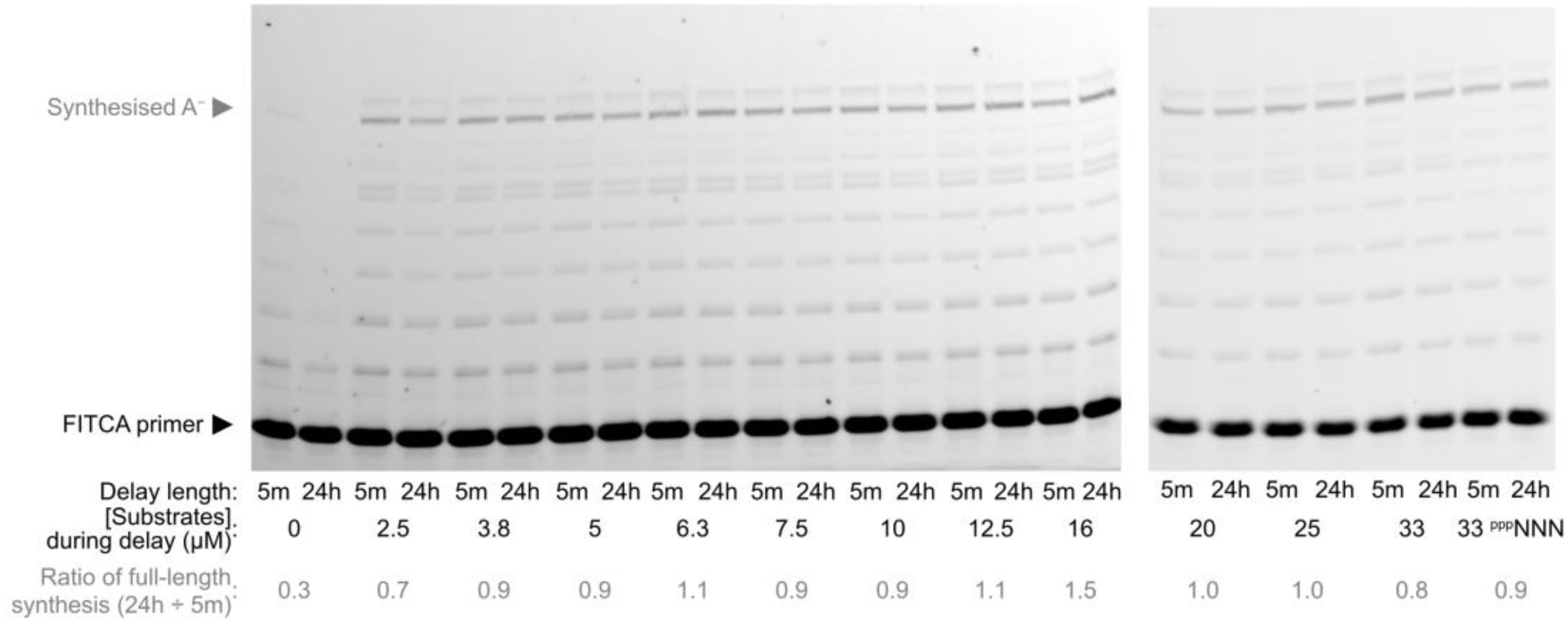
Primer extension upon acid-denatured duplex, following delayed addition of varying amounts of substrates after neutralisation. Reactions were set up as in Fig. 1c, but with different concentrations of triplet present upon neutralisation and freezing (shown are estimated eutectic phase triplet concentrations after neutralisation). After a delay of 5 minutes or 24 h, ribozyme was added (alongside remaining substrates to a final eutectic phase concentration of 25 µM) before continued incubation at −7°C. The proportional reduction in full-length product signal with a longer delay in each pair of lanes (each triplet concentration) is calculated by gel densitometry. Little reannealing appears to occur even down to low triplet concentrations. ^ppp^NNN: Instead of just the triplet substrates in Fig. 1a, all 64 triplet sequences were added as substrates; the strand coating effect was maintained.

**Fig. S4.**
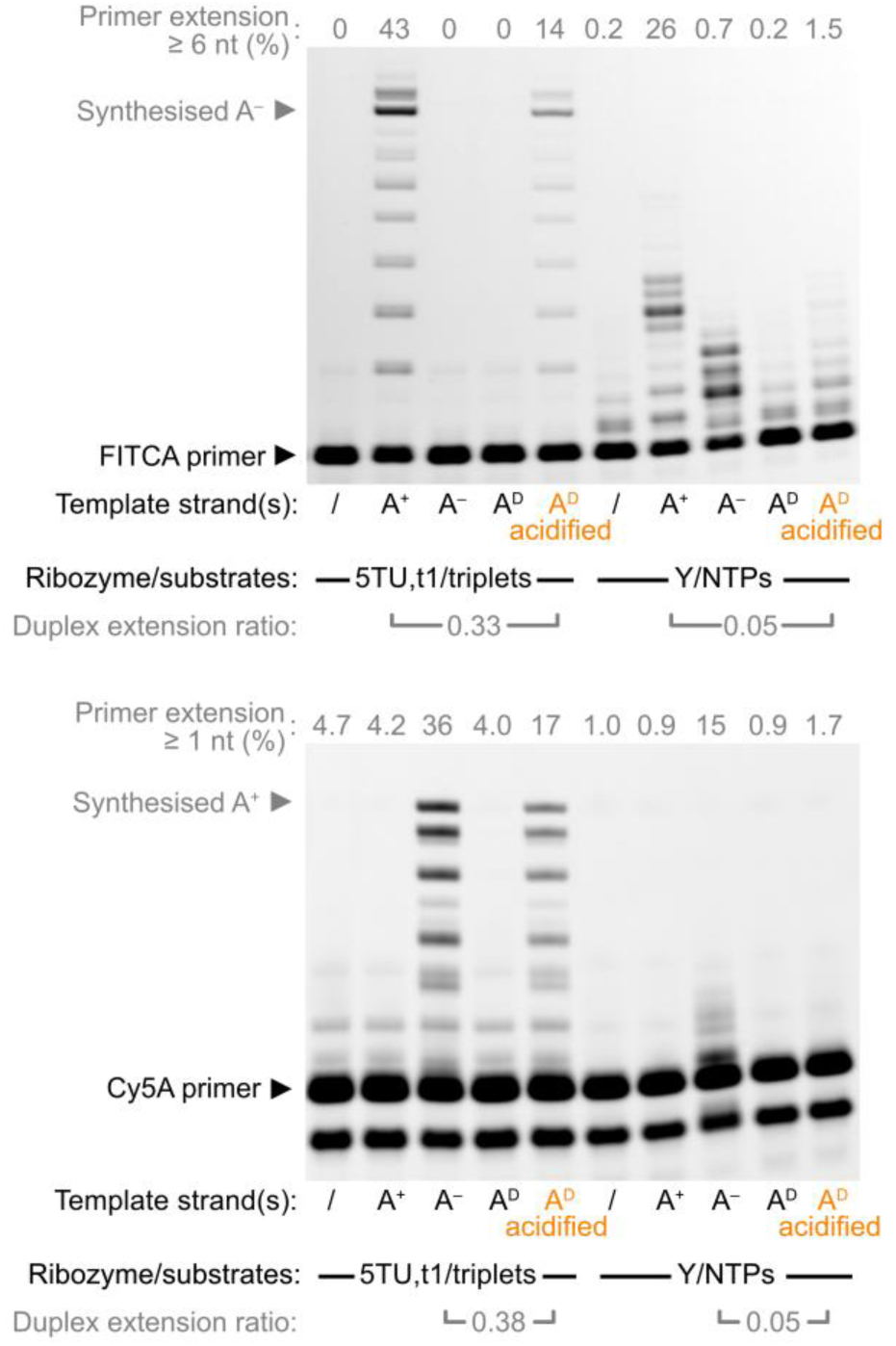
Primer extension activity of triplet polymerase versus monomer polymerase ribozymes using RNA duplex templates. Denaturing PAGE of primers (0.1 µM each FITCA (imaged above) and Cy5A (imaged below)) extended upon different A templates (0.1 µM) in extension buffer (100 mM MgCl_2_, 50 mM Tris·HCl pH 8.3, 0.05% Tween-20). This was catalysed by (left) 0.5 µM TPR (with 2.5 µM of each triplet in the Fig. 1a synthesis scheme, −7°C 2 days), or (right) using 0.5 µM mononucleotide polymerase ribozyme Y^24^ (with 0.5 mM of each of the four nucleoside triphosphates (NTPs), −7°C 12 days). Yields of primer extension (≥ 6 nt (upper panel) or ≥ 1 nt (lower panel)) in each lane were estimated by gel densitometry. Proportionally less extension was achieved by acidification of duplex when using mononucleotides as substrates compared to triplets, as judged by the duplex extension ratio on each template strand, calculated using estimates of ‘% extension above background’ from densitometry: (‘A^D^ acidified’ ‒ ‘/’) ÷ (‘A^+ or‒^’ ‒ ‘/’)

**Fig. S5.**
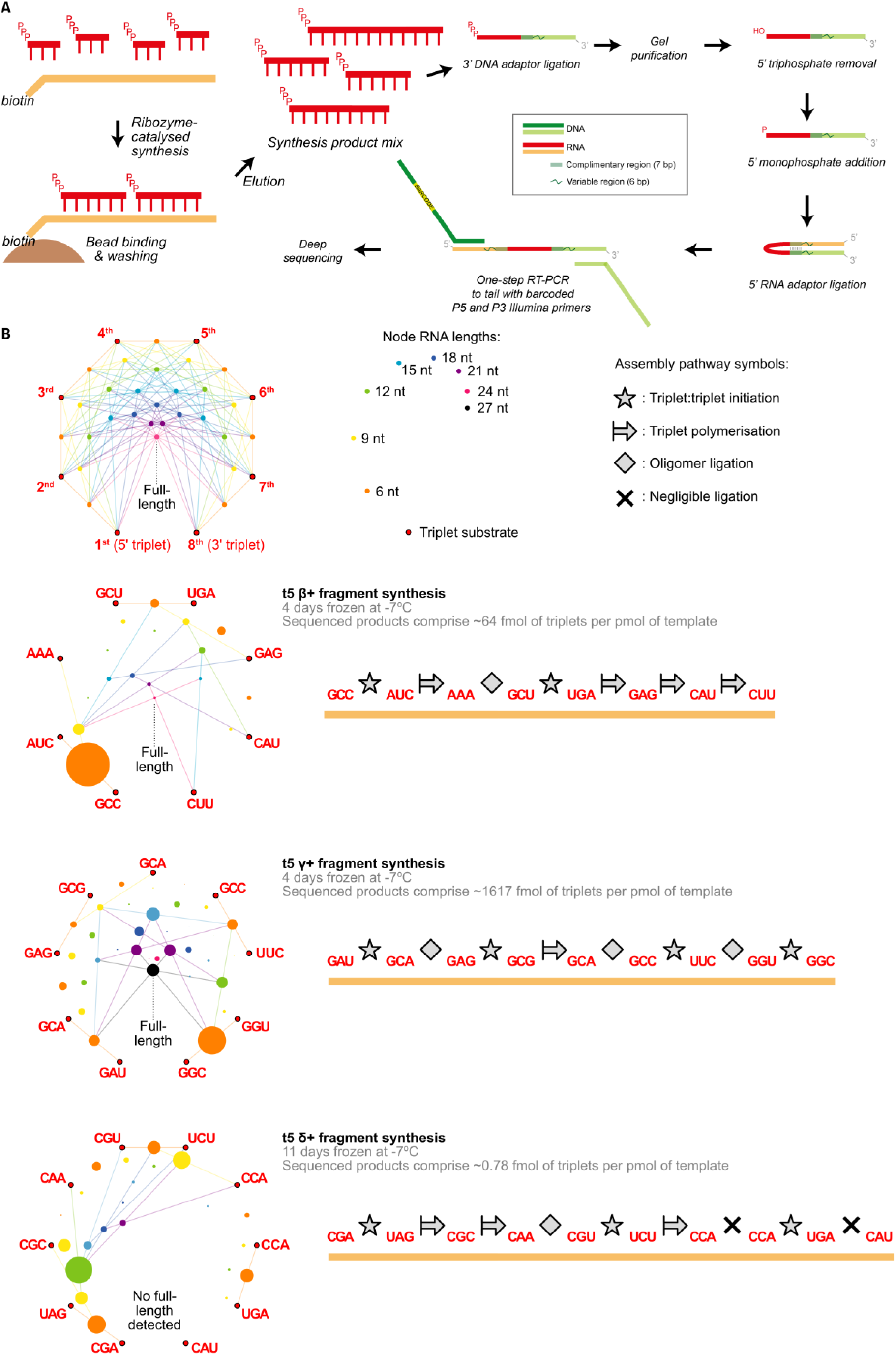
Primerless synthesis pathways of triplet polymerase ribozyme fragments. (**a**) Sequencing scheme for identification of synthetic intermediates during templated synthesis of ribozyme fragments using template-complementary triplets alone. (**b**) Mapping of synthesized products and identification of potential synthesis pathways. (Above) Map of theoretical pathways available for synthesis of a 24 nt RNA from eight triplet substrates. Each node represents a synthesis intermediate, of increasing length towards the center of the map. The intermediate at each node can be synthesized from the intermediates or triplets at each end of a line passing through the node. Larger intermediates can potentially be generated via multiple routes. (Below left) Maps of products detected during primerless synthesis of three fragments of the t5 ribozyme^4^: β+ (top), γ+ (middle) and δ+ (bottom). These syntheses proceeded with very different efficiencies, and the estimated overall product yields are noted in triplet equivalents (γ+ > β+ > δ+). The maps themselves display the distribution of polymerised triplets: the area of each node is proportional to the abundance of that intermediate multiplied by its length. Node sizes are uniformly scaled in each map such that the sum of intermediate node areas is constant; the triplet substrate nodes (not sequenced) are sized arbitrarily. On each map, lines are drawn to represent putative synthesis pathways based upon intermediate abundances; other pathways are technically possible but may reflect intermediate distribution less well. (Below right) Synthesis pathways of the three fragments. The order of triplet substrates is displayed in a 5’ to 3’ direction with intervening symbols indicating how they are joined together into RNA products. The symbols chosen reflect the lines drawn on the maps to the left – indicating sites of initiation (triplet:triplet ligation), polymerisation (triplet addition to the 3’ of an oligonucleotide) or instances of ligation (of oligomer substrates larger than triplets). The β fragment exhibits two sites of initiation, followed by polymerisation from each with their concomitant ligation on the path to the full-length fragment. The γ fragment synthesis is characterized by multiple sites of initiation and independent ligation of the resulting oligomers; there is evidence of further initiation sites in the 5’ section of the fragment beyond those displayed in the synthesis scheme. Some junctions do not appear to exploit polymerisation (i.e addition of only a triplet) and use longer substrates. This primerless assembly proceeded efficiently, perhaps reflecting the availability of multiple initiation sites. The δ primerless synthesis was inefficient, with negligible ligation at two junctions, and as a result no full-length product was detected. Those products that were observed suggested a pattern of two initiation sites, followed by polymerisation and ligation. The triplet:triplet ligation observed across the three templates is evidence of widespread triplet substrate coating of single-stranded templates.

**Fig. S6.**
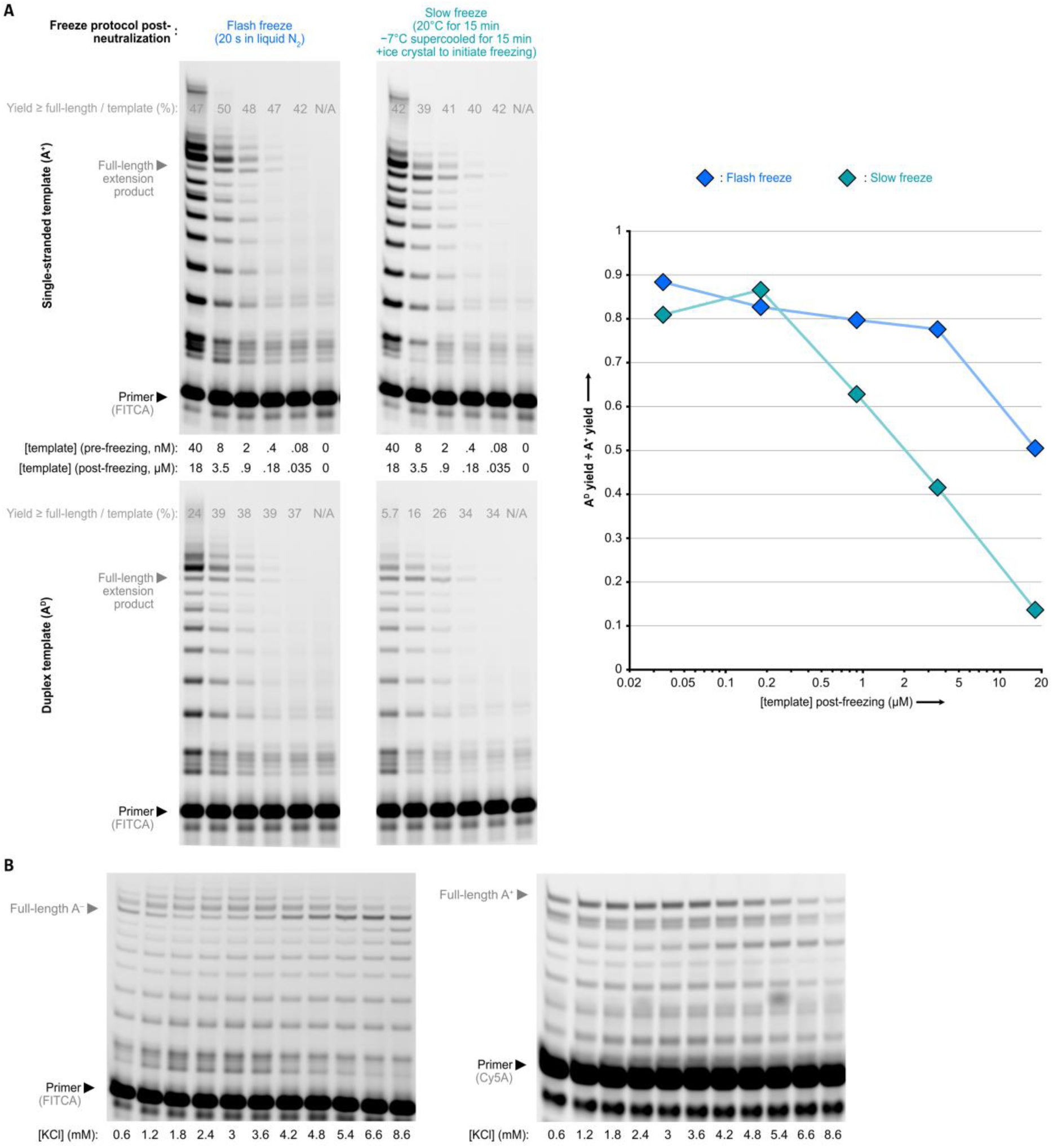
Experimental constraints on the freeze/thaw/pH cycling protocol. (**a**) Flash-freezing maximises polymerisation efficiency at high RNA duplex concentrations. A single cycle of TPR-catalysed replication (as in Fig. 2b, with 2.4 mM KCl, −7°C for 40 h) was applied to different concentrations of single-stranded RNA (top) or duplex RNA (bottom). Gel densitometry after PAGE separation of products (left) allowed calculation of the proportion of primers extended to the full length of the template in each reaction; this was divided by the template concentration to obtain the yield per template. When using single-stranded templates, this yield was broadly constant. However, the yield dropped when using high concentrations of duplex templates. This was dependent on the protocol used to freeze the reaction after acidification, heating and neutralisation: a flash-freeze in liquid N2 over 20 seconds (blue) maintained yield at a higher concentration of duplex than a slow freeze over half an hour (teal, 15 min at room temperature, 15 min at −7°C supercooled, then frozen by ice crystal addition). Right, plot of yield ratio between duplexes and single-stranded RNAs across the template concentrations tested for each freeze protocol. This decline likely reflects reannealing of the two strands of a duplex before freezing establishes a high enough concentration of triplet substrates to coat the strands and block reannealing; this is more severe at higher duplex concentrations and longer times between neutralisation and eutectic phase formation. In effect, these parameters govern the relative weights of the ‘reannealing’ and ‘strand coating’ arrows in Fig. 1e and thus the productivity of the replication cycling. Efficient duplex use (∼80% vs. ssRNA) was nonetheless observed at eutectic phase concentrations of 3.5 µM (after flash-freeze) or 0.8 µM (after slow freeze). This concentration is a critical parameter of any cycling protocol as it governs the maximum concentration of each RNA species that can be generated via replication. Reassuringly, ribozymes are operational at these levels, including the TPR itself (Fig. S1b). Therefore, sufficient catalysts could theoretically accumulate via this cycling protocol to drive biological processes. (**b**) Accumulation of KCl during cycling moderately inhibits extension on template A. Shown are single cycles of replication with varying concentrations of KCl, but otherwise set up as in Fig. 2b, with 4 nM starting duplex A^D^. The [KCl] shown includes the 0.6 mM increase resulting from the starting cycle of acidification and neutralisation. Freezing likely yields molar ionic concentrations in the eutectic phase, and at the highest [KCl], less ice cystal growth may occur as eutectic composition is reached earlier – effectively diluting other reaction components as a result and changing the pattern of primer extension. To attenuate this, iterative cycling underwent occasional dilution with fresh reaction mix lacking KCl.

**Fig. S7.**
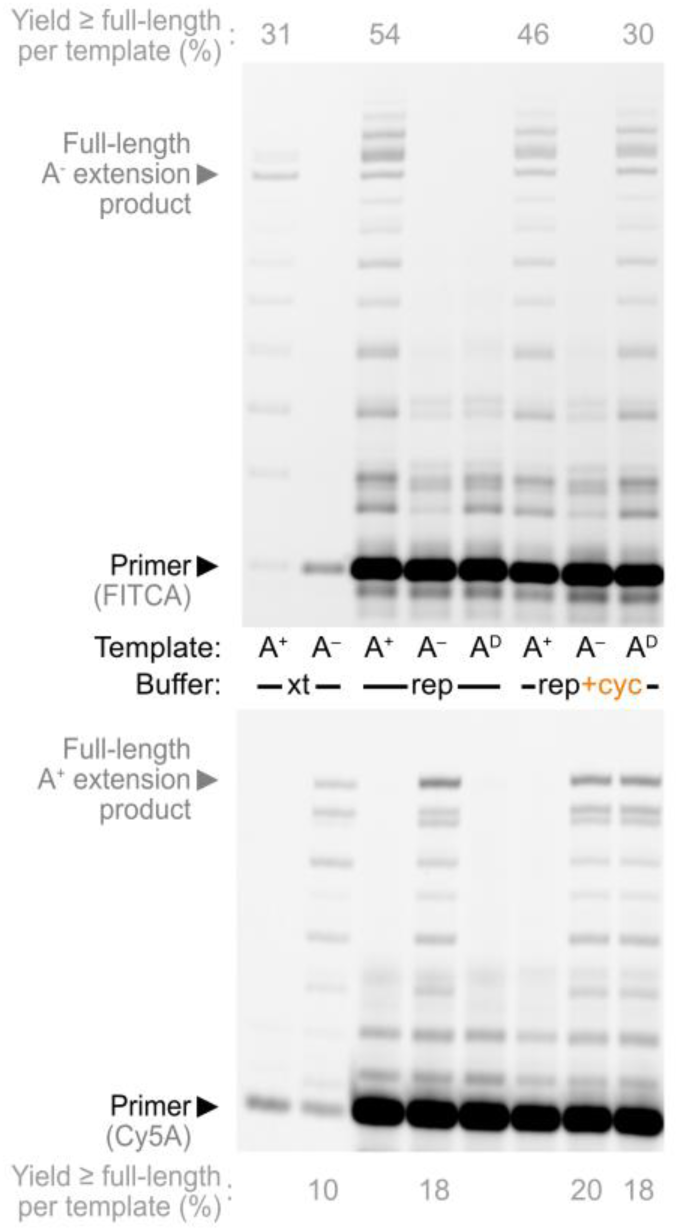
Primer extension efficiency upon single-stranded and duplex templates in extension buffer and replication buffer. Strand synthesis in replication buffer matches that in extension buffer (in which the ribozyme was originally evolved^4^); implementation of a pH cycle has little effect on subsequent extension, but efficiently unlocks the templating activity of RNA duplex. Extension buffer (xt): 100 mM MgCl_2_, 50 mM Tris·HCl pH 8.3 @ 25°C, 0.05% Tween-20, 0.1 µM of each primer and 2.5 µM of each triplet substrate in the scheme of Fig. 1a, 0.5 µM TPR, 0.1 µM f.c. template; −7°C frozen 48 h, ∼10-fold solute concentration in the eutectic phase^22^. Replication buffer (rep): 0.4 mM MgCl_2_, 2.4 mM KCl, 1 mM CHES·KOH pH 9.0 @ 25°C, 0.01% Tween-20, 0.1 µM of each primer and 0.1 µM of each triplet substrate in the scheme of Fig. 1a, 20 nM TPR, 4 nM f.c. template; −7°C frozen 48 h, ∼440-fold solute concentration in the eutectic phase. Replication buffer after cycling (rep+cyc): as rep but with addition of 0.6 mM HCl, incubation at 80°C for 2min, and addition of 0.6 mM KOH before flash-freezing and incubation. Densitometry gives estimates of full-length product yield per template.

**Fig. S8.**
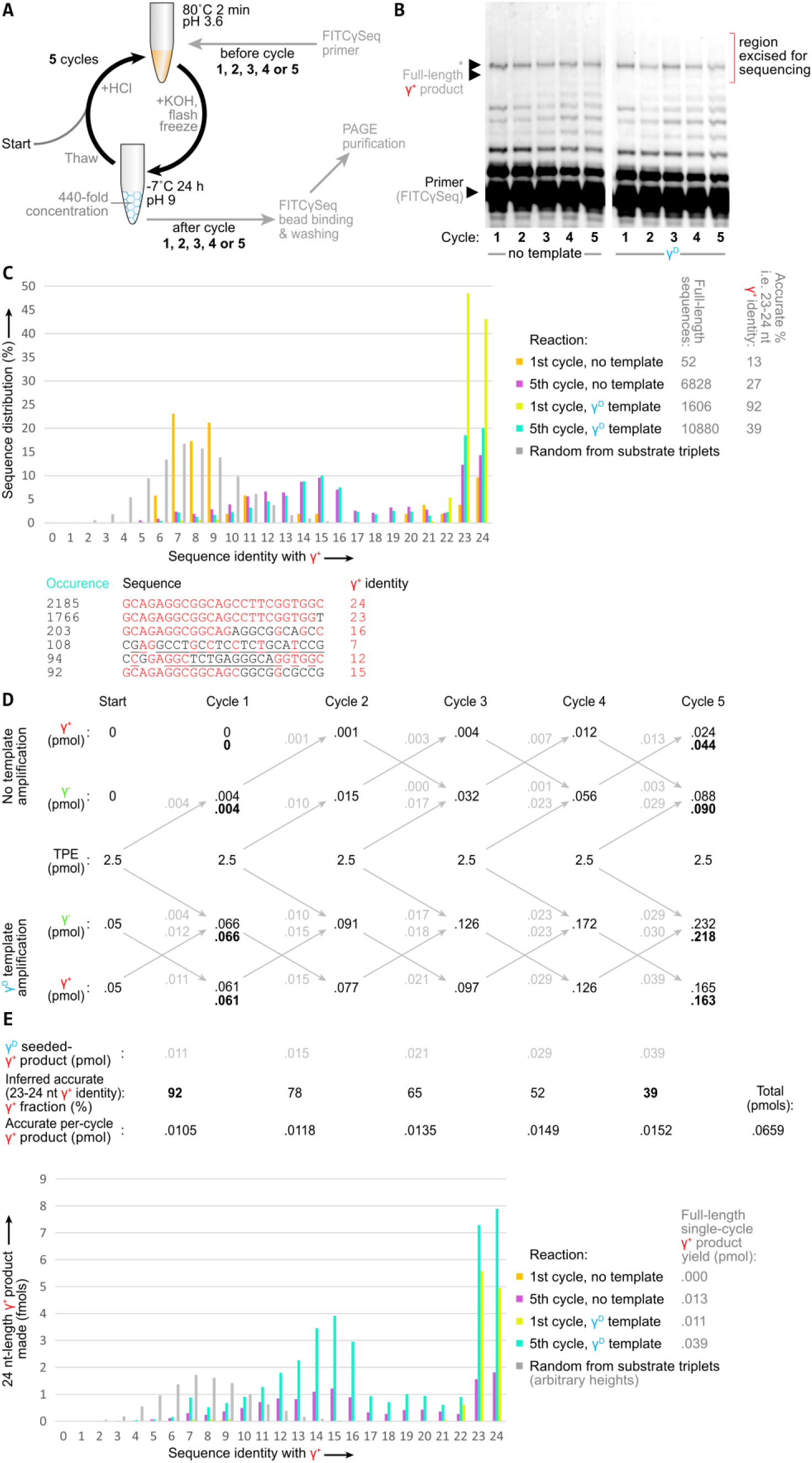
Sequencing and analysis of γ fragment replication reactions. (**a**) Scheme for cycle-specific sequencing of replication products. Reactions were set up as in Fig. 4d, but with 0.1 µM FITCγSeq sequencing primer added during acidification before one of the 1^st^ to 5^th^ cycles. Reactions were then stopped after this single extra cycle of sequencing primer extension. These primers were then separated from the other FITCγ primers via a biotin moiety on FITCγSeq, obtaining a cycle-specific record of primer extension. Iterative cycling with FITCγSeq alone was not performed as the lengthened 5’ of this RNA (needed to provide an rtPCR primer binding site of sufficient length for sequencing) reduced replication yields. (**b**) Denaturing PAGE separation of FITCγSeq primer extensions in (a). The indicated gel regions (containing full-length products) for cycles 1 and 5 in γ^D^-seeded and no template reactions were excised and the primers therein ligated to 3’ adapters, amplified by rtPCR, and subjected to high-throughput sequencing. The major band present in this region (*) is a template-independent ligation artefact seen with the longer sequencing primer; only full-length +24 nt products were further analysed. (**c**) Proportion of accurate replication products. The indicated number of such 24 nt length sequences were obtained from each sequencing reaction. These are plotted according to their degree of sequence identity with the expected γ^+^ sequence. Also plotted is a distribution of sequence identity amongst 24 nt sequences randomly assembled from the substrate triplets present in the reaction. Three classes of product are apparent. A single cycle of replication on γ^D^ generates accurate full-length γ^+^ sequences (23-24 nt identity). Further cycles also generate products with partial identity (13-16 nt identity), potentially derived from template-switched incomplete extension products. Finally, there are sequences with no discernible identity with γ^+^ (7-12 nt identity), dominating the output of a single cycle without template, where no γ^-^ sequence is present to direct γ^+^ formation. The six most common individual sequences in the 5-cycle γ^D^ replication sample are shown below the chart; interestingly, the two of these sequences with very low γ^+^ identity show complementarity (underlined) to parts of the type 1 ribozyme (see Fig. S1a), a spontaneous instance of TPR acting as a template. (**d**) Model of replication product synthesis yields. To compare the levels of sequence class synthesis between different reactions, we needed to adjust the sequence distribution by the observed yields of full-length product in each cycle. Unfortunately the artifact band (*) prevented quantification of FITCγSeq extension to full-length product in (b). We therefore modelled full-length product accumulation during the five cycles of seeded and unseeded replications - templated from product strands, TPR, and degraded TPR. For amplification constants, we used the γ^D^ templating efficiencies observed in the first cycle in Fig. 4d (0.228-fold γ^+^ from γ^-^, 0.239-fold γ^-^ from γ^+^, and 0.00167 γ^-^ from each TPR per cycle). An increase in the latter parameter of 0.0025 per cycle (to account for the increased templating capacity of degraded TPR (Fig. 4d) that accumulates in the polymerisation steps) gave a reasonable match of the modelled yields to the observed yields (in bold, Fig. 4d) after five cycles. (**e**) Per-cycle yields of accurate γ^+^ product formation. The percentage of accurate products in each cycle was estimated by interpolating the observed accuracies in the first and fifth cycles. This was then multiplied by the modelled per-cycle yields of all γ^+^ products, giving an estimated synthesis yield of 0.065 pmols accurate γ^+^ strand during five cycles of replication of 0.05 pmols γ^D^. This yield does not include synthesis products exceeding full-length. Below, the sequence product distribution in (c) for each sample was scaled by the modelled yield of all γ^+^ products in the corresponding cycle.

**Fig. S9.**
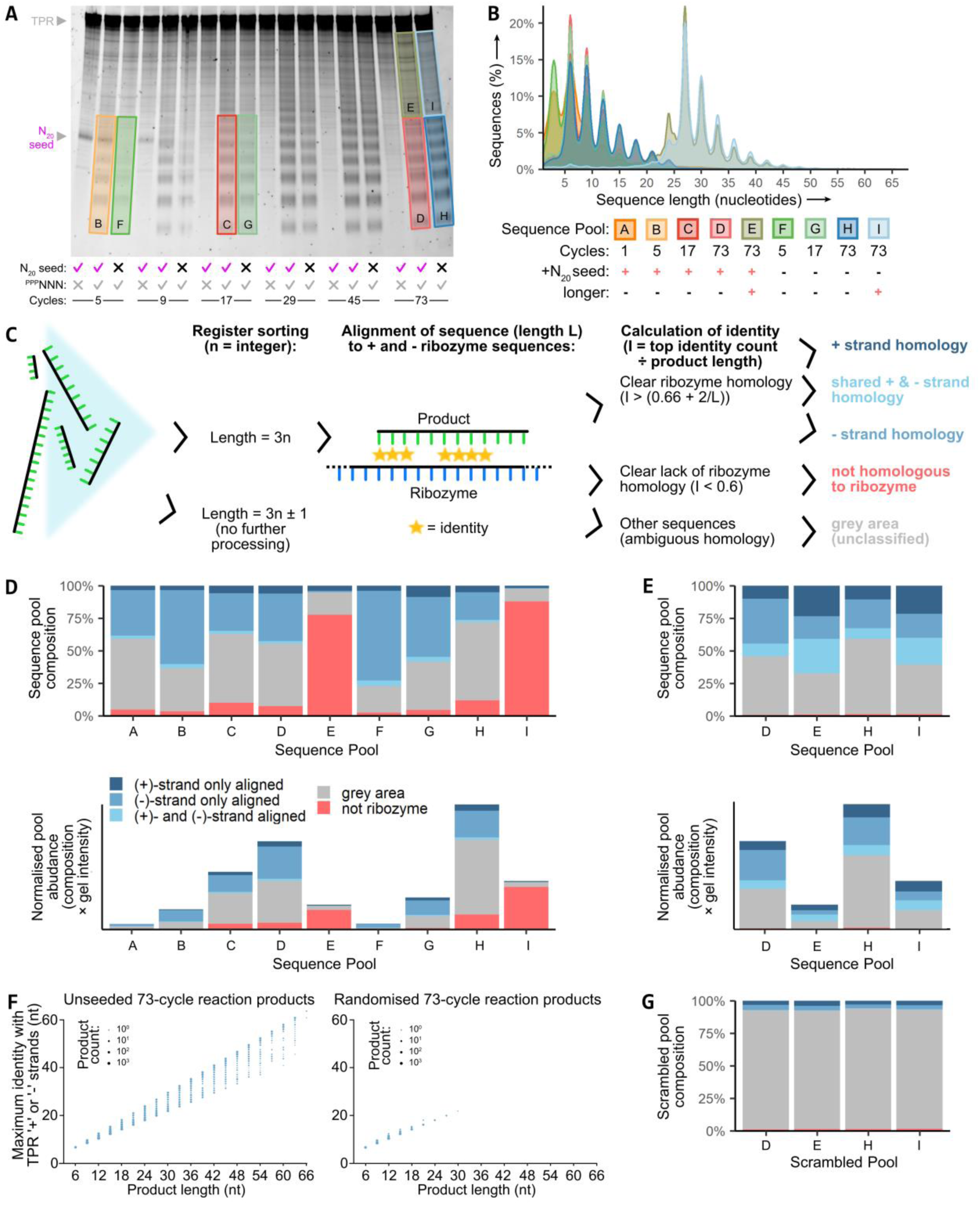
Sequencing and classification of amplification products made from ^ppp^NNN substrates. (**a**) Isolation of amplification products. We excised the indicated regions of the gel used to PAGE separate ^ppp^NNN amplification products (Fig. 5b) and eluted and precipitated the RNAs therein. To sequence these products we used a variant of the protocol in Fig. S5a. There, alkaline phosphatase and polynucleotide kinase treatments generated 5’ monophosphates to enable 5’ adaptor ligation; here, RNAs were instead treated with a pyrophosphohydrolase to selectively convert 5’ triphosphates to monophosphates (to promote selective ligation and sequencing of ribozyme-synthesised products; nonetheless, a low level of TPR molecules/fragments were sequenced – see (f)). The named pools A-I sampled different stages of the N_20_ seeded and unseeded amplifications with ^ppp^NNN; the N_20_ seed itself, lacking a 5’ triphosphate group, would not be sequenced. Pool A was derived from excising the corresponding 9-27 nt region of a separate equivalent 1-cycle seeded amplification. (**b**) Length distributions observed within the pools of sequenced RNAs. The strong triplet register bias confirms sequencing of ribozyme-synthesised RNAs. (**c**) Workflow for classification of the sequenced products. Only triplet-register products were analysed. Amplification products that exceeded stringent length-dependent sequence identity thresholds at any point when aligned along the (+) strand sequences of either 5TU or t1 TPR subunits (or their (−) strand complements) were classed as possessing ribozyme homology. These were further subdivided by the (+) or (−) strand they were matched to; some showed homology to both (unsurprising in a hairpin-rich RNA with internal complementarity) and were classed separately. Sequences that clearly did not align to the ribozymes or their complements at any point were also classed separately. Sequences between the indicated identity thresholds could not be easily categorised and were excluded from further analysis. (**d**) Levels of ribozyme homology within RNA product pools. Top: the fractions of each sequenced product pool that showed ribozyme homology as determined in part (c). A substantial fraction exhibits complementarity to ribozyme, though this decreases after many cycles. Bottom: absolute amounts of different product sequences. Here, total column heights are proportional to amounts of synthesised and triplet register RNA, based upon intercalator fluorescence in the excised region in part (a); columns are subdivided as in the top chart. All categories of product increase in absolute abundance over the course of cycling. (**e**) Reclassification of longer amplification products. Most of the > 27 nt sequences apparently exhibited no overall ribozyme homology (see part (d)). However, this was partly an artefact of aligning all along these longer sequences. Here, only the 4^th^-12^th^ nt of each sequence was aligned, revealing ribozyme homology in a similar proportion of longer sequences (pools E & I) to those in the shorter fractions (pools D & H). Thus a substantial fraction of the longer amplification products exhibit local but not global ribozyme homology; they may have been generated by recombination or seeded from ribozyme-derived shorter sequences. Note that the 9 nt length window used here is too short to definitively class sequences as ‘not derived from ribozyme’ using our identity thresholds. (**f**) TPR identity amongst sequences classed as (+)-strand homologous. Shown are levels of identity amongst sequences meeting the identity threshold for (+) or (+) & (−) strand homology (with other sequences plotted in Fig. 5e), for both sequenced synthesis products from the unseeded 73-cycle reaction (left), and for a simulated pool of randomised RNAs of identical composition (right). The reaction products contain a population of sequences with complete or near-complete (+)-strand identity, exhibiting a reasonably uniform length distribution. These likely derive from background sequencing of ribozyme (or degradation products thereof, despite requiring a 5’ phosphate or triphosphate for recovery). A second population of products (<30 nt, and more abundant than in a pool of randomised sequences) has only partial (+) strand homology indicating a synthetic origin. (**g**) Negligible ribozyme homology is observed amongst scrambled pool sequences. 9 nt sequences, generated from random triplet assortments (matching the triplet compositions of the indicated pools) were classified as in part (c).

**Fig. S10.**
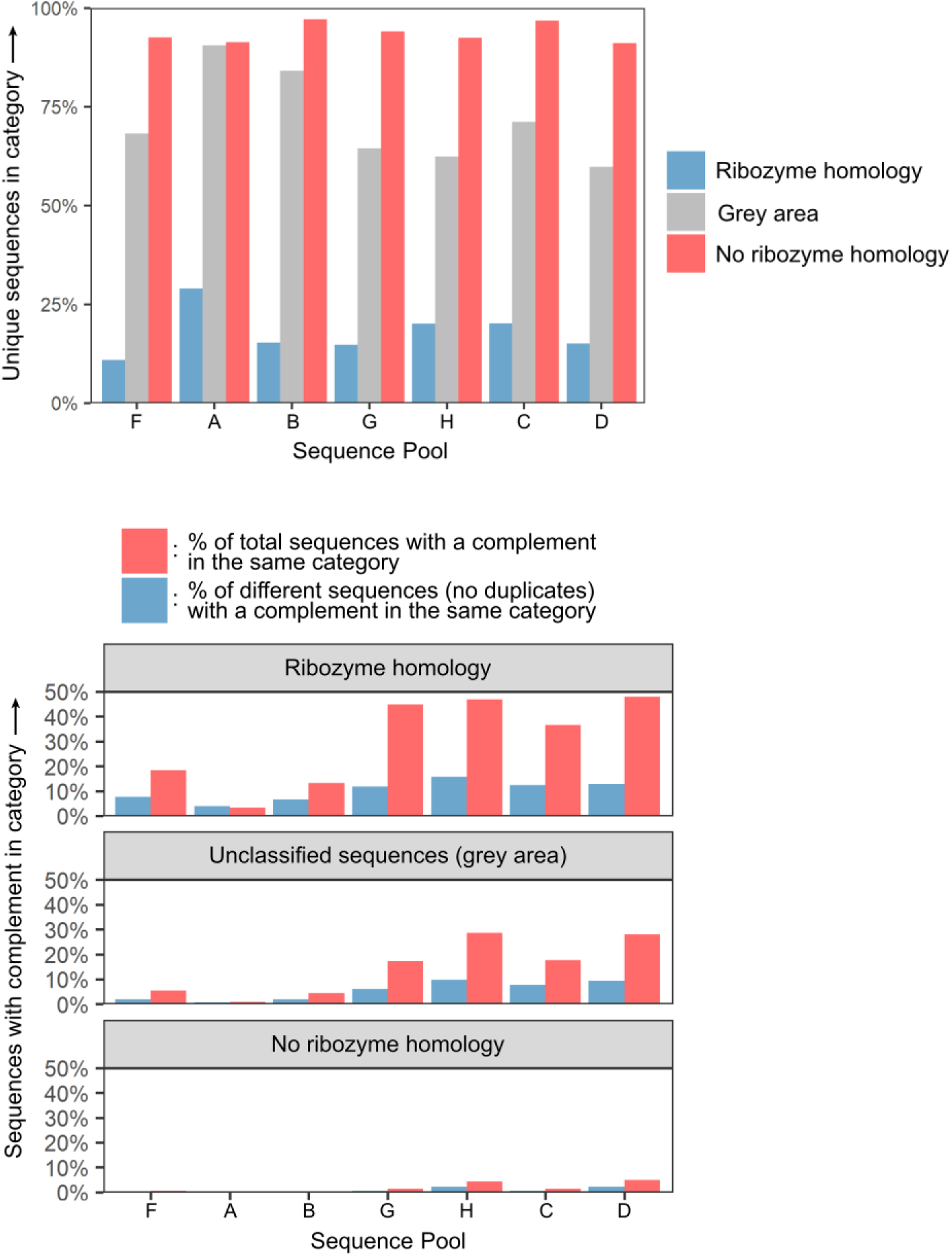
Relationships of RNA amplification products to one another. Top: the percentages of each 9-27 nt sequenced replication product pool subcategory (classified by ribozyme homology) that comprise unique sequences. Across all samples, the majority of sequences with homology to ribozyme were present multiple times in the sequence data; the majority of those with no evidence of homology were unique. Pools labelled as in Fig. S9. Bottom: Prevalence of complementary sequences among amplification reaction 9-27 nt products (from Fig. S9), analysed by subcategory. This is expressed as the percentage of sequences in each replication product pool subcategory (classified by ribozyme homology) with a perfect complement in the same subcategory (red). This calculation was also performed after discarding repeat sequences in each subcategory, showing the percentage of different sequences with a complement in that subcategory (blue). The sequences unrelated to ribozyme showed little complementarity with each other, making reciprocal primerless replication an unlikely mechanism for their amplification. There was some complementarity among ribozyme-related (‘+’ and ‘-’) sequences, though some may arise from the presence of regions of internal complementarity within each ribozyme.

**Supplementary Table 1.**
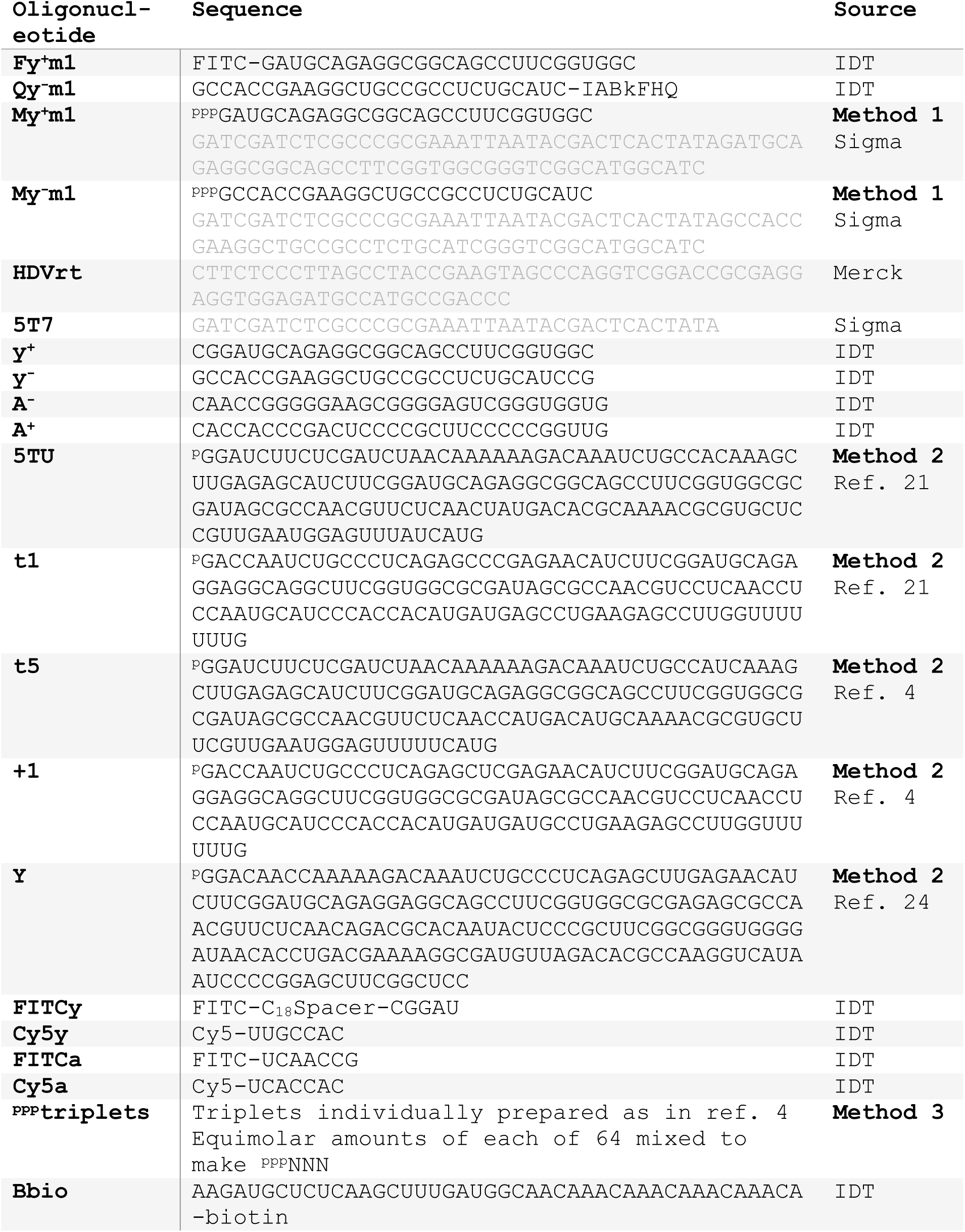
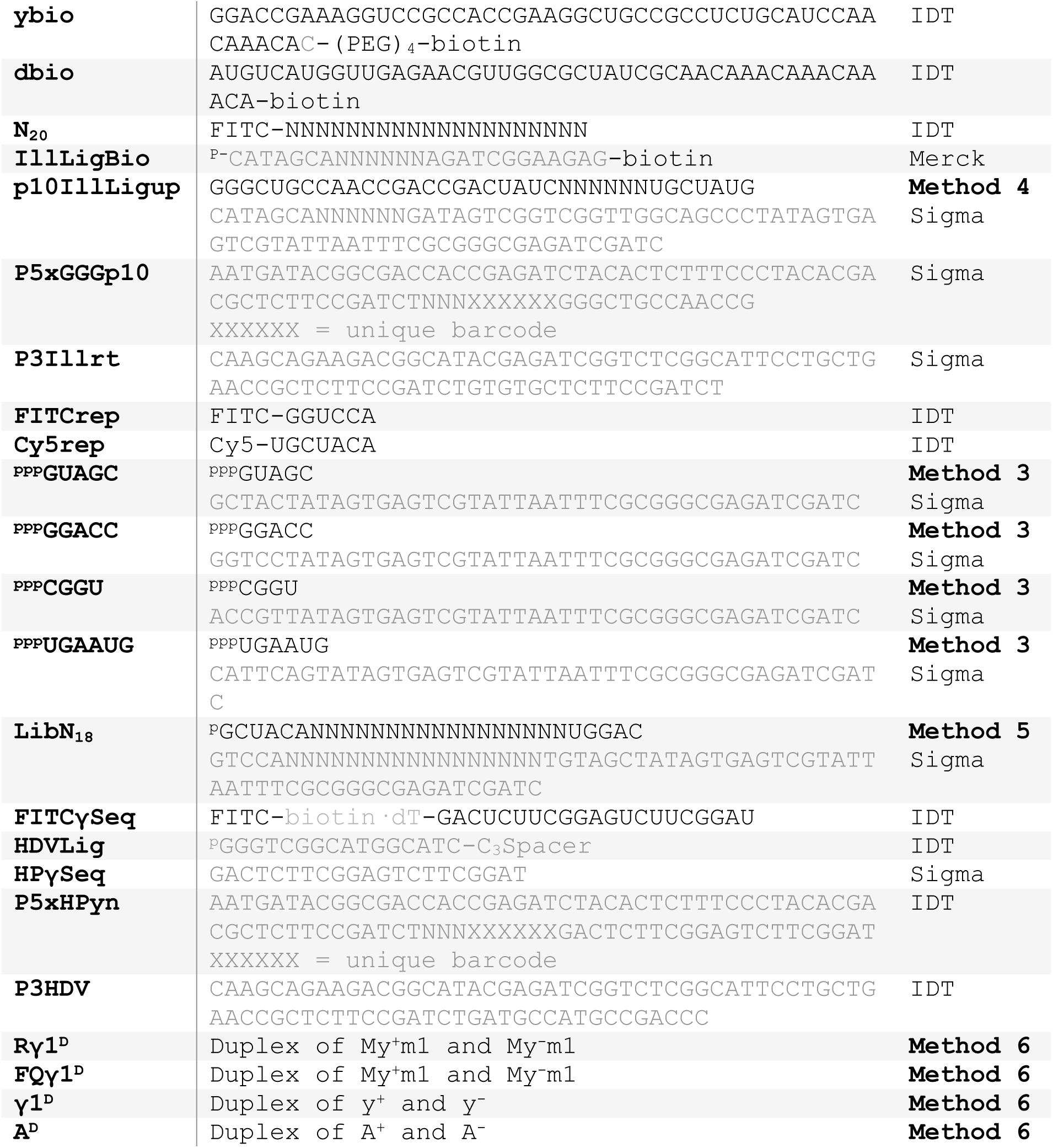
– Oligonucleotide sequences. Oligonucleotide sequences are written 5’ to 3’ below; DNA sequences are in grey, RNA sequences are in black. RNAs synthesised from DNA sequences were prepared by methods 1-6 as described in the Methods section. All oligonucleotides (except DNAs used to make transcription templates) were PAGE-purified before use and concentrations calculated using a Nanodrop spectrophotometer together with sequence-specific 260 nm extinction coefficients derived using OligoCalc^40^.

**Supplementary Table 2.** – Custom scripts. Custom scripts will be made available at GitHub. In scripts from ‘Sequence_lengths’ onwards, datasets ’RZ5N’, ’N17L’, ’N73L’, ’N73H’, ’F201N’, ’F205N’, ’FN17L’, ’FN73L’, and ’FN73H’ correspond to samples A-I respectively (Fig. S9).

| **Name** | **Language** | **Description** | **Lines** | **Dependencies** |
| --- | --- | --- | --- | --- |
| **specify** | Python 3 | Generate .txt file of all possible in frame species from a template. | 25 | NumPy, Biopython 1.71<sup>41</sup> |
| **intermas**<br>**ter1.2** | Python 3 | Map clipped HTS reads to list of expected species, with option to tolerate errors. | 98 | NumPy, Biopython 1.71<sup>41</sup> |
| **tally** | Python 3 | Measures and counts sequence identity with expected synthesis product | 29 |  |
| **24ntrand**<br>**om** | Python 3 | Generates random sequences from a defined set of triplets | 31 |  |
| **Sequence**<br>**_lengths** | Python 3 | Prepares sequences to plot lengths of sequences (inputs from .fasta files) | 59 | pandas, os, Biopython |
| **Plot_seq**<br>**uence_le**<br>**ngths** | R | Plots lengths of sequences before filtering (inputs from Sequence_lengths.py) | 50 | ggplot2, tidyverse, tidyr, dplyr, RColorBrewer |
| **Alignmen**<br>**ts** | Python 3<br>(Jupyter<br>notebook<br>) | Aligns sequences to ribozyme sequences after filtering, assess uniqueness, complementarity and nucleotide composition (inputs from .fasta files) | 1317 | pandas, os, Biopython, numpy |
| **Family_b**<br>**ox_codon**<br>**s** | Python3 | Assess continuity of family box codons in pool H+I and random sequences (inputs from .fasta files) | 333 | pandas, os, Biopython |
| **Plot_ali**<br>**gnment_c**<br>**ount** | R | Plots of the bins of different alignments (inputs from Alignments.ipynb) | 231 | ggplot2, tidyverse, tidyr, dplyr, RColorBrewer |
| **Plot_ali**<br>**gnment_r**<br>**ibozyme** | R | Plots scores of aligned sequences and their position on the ribozyme (inputs from Alignments.ipynb) | 55 | ggplot2, tidyverse, tidyr, dplyr, RColorBrewer |
| **Plot_fam**<br>**ily_box_**<br>**codons** | R | Plots percentage of sequences with contiguous family box codons (input from Family_box_codons.py) | 47 | ggplot2, tidyverse, tidyr, dplyr, RColorBrewer |
| **Plot_uni**<br>**queness** | R | Plots uniqueness of sequences in different | 48 | ggplot2, tidyverse, |
|  |  | categories (input from Alignments.ipynb) |  | tidyr,<br>dplyr,<br>RColorBrewer |
| **Plot_complementarity** | R | Plots complementarity of sequences in different categories (input from Alignments.ipynb) | 57 | ggplot2,<br>tidyverse,<br>tidyr,<br>dplyr,<br>RColorBrewer |
| **Plot_identity_vs_length** | R | Plots identity of sequences with the ribozyme of different length (input from Alignments.ipynb and Intensities_HI.csv) | 231 | ggplot2,<br>tidyverse,<br>tidyr,<br>dplyr,<br>RColorBrewer<br>, plyr |
| **Plot_nucleotide_composition** | R | Plots nucleotide composition of triplets over multiple cycles in non-ribozyme sequences (input from Alignments.ipynb) | 72 | ggplot2,<br>tidyverse,<br>tidyr,<br>dplyr,<br>RColorBrewer |

**Supplementary Table 3.** – Codon usage in amplification products. We calculated the triplet composition within pool H sequences (Fig. S9) from the 73-cycle unseeded amplification reaction to understand the partition of triplets between substrate and products in the analytical sample. The fraction of the sequences constituting each triplet was then multiplied by the observed yield of RNA product in the reaction (the equivalent of 284.5 pmols of triplets; see Fig. 5c) to estimate the consumption (%) of each triplet (from the 12.5 pmols of each of the 64 triplets available in the analytical sample) shown to the right of each triplet sequence. Some triplets’ consumptions are calculated to exceed 100%; this may reflect slight sequence/structure/length preferences of RNA intercalation, adapter ligation and/or rtPCR in the work-up, frequent when sequencing short RNAs^39^. GC-rich triplets are particularly depleted, and their high degree of utilisation might also reflect their better capacity to initiate RNA synthesis on the growing product pool (Fig. S5).

|  |  |  |  |  |  |  |  |
| --- | --- | --- | --- | --- | --- | --- | --- |
| UUU | 1 | UCU | 10 | UAU | 3 | UGU | 5 |
| UUC | 2 | UCC | 16 | UAC | 2 | UGC | 30 |
| UUA | 0 | UCA | 7 | UAA | 0 | UGA | 10 |
| UUG | 7 | UCG | 18 | UAG | 2 | UGG | 34 |
| CUU | 8 | CCU | 83 | CAU | 22 | CGU | 27 |
| CUC | 40 | CCC | 129 | CAC | 21 | CGC | 79 |
| CUA | 8 | CCA | 75 | CAA | 5 | CGA | 33 |
| CUG | 78 | CCG | 143 | CAG | 29 | CGG | 102 |
| AUU | 3 | ACU | 3 | AAU | 0 | AGU | 3 |
| AUC | 6 | ACC | 16 | AAC | 2 | AGC | 13 |
| AUA | 4 | ACA | 3 | AAA | 0 | AGA | 7 |
| AUG | 9 | ACG | 8 | AAG | 10 | AGG | 47 |
| GUU | 17 | GCU | 80 | GAU | 51 | GGU | 72 |
| GUC | 35 | GCC | 85 | GAC | 31 | GGC | 60 |
| GUA | 17 | GCA | 161 | GAA | 21 | GGA | 103 |
| GUG | 114 | GCG | 84 | GAG | 93 | GGG | 88 |

**Supplementary Table 4.**
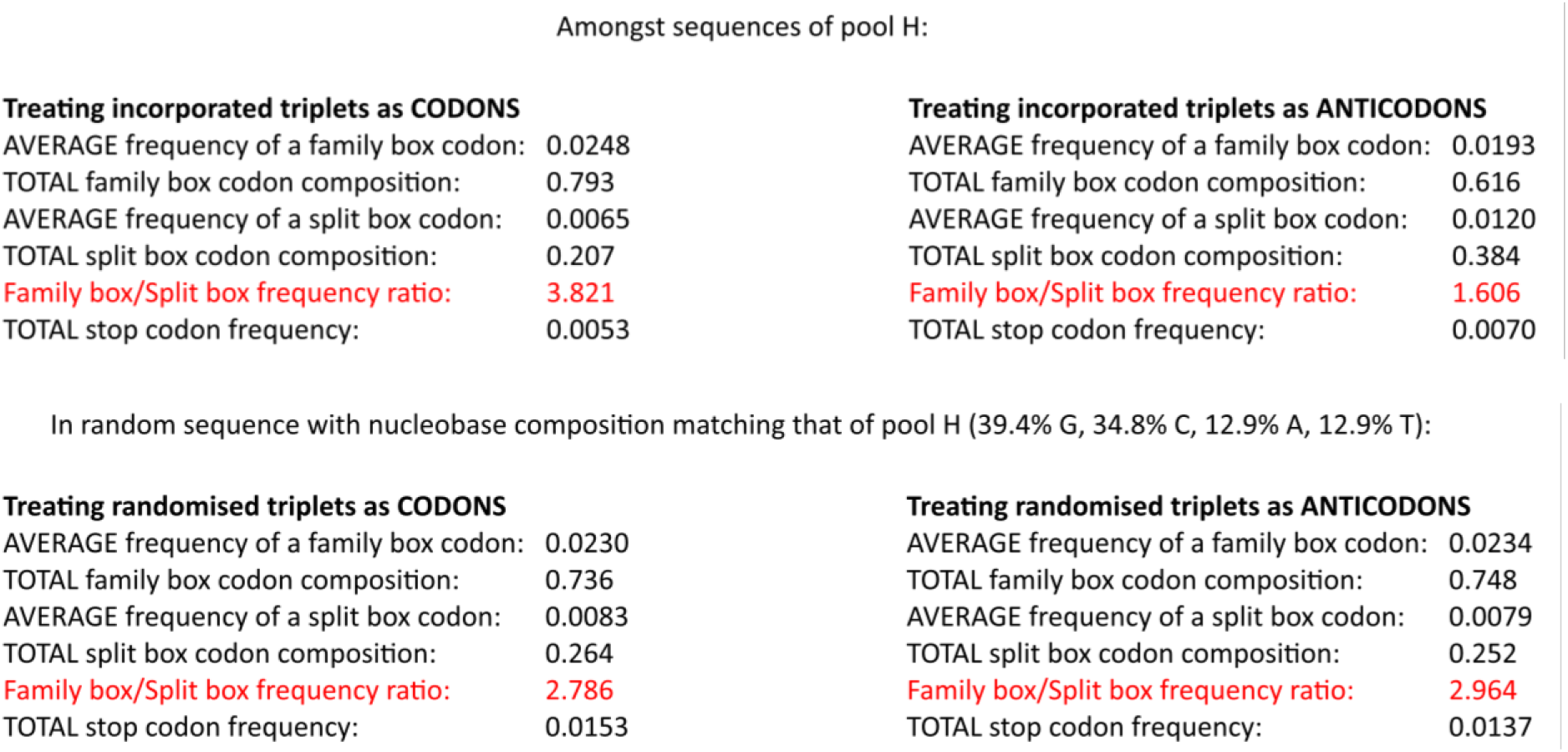
– Codon class enrichment in amplification products. Comparison of ribozyme-generated product composition with potential early genetic code composition. The triplets highlighted in Supplementary Table 3 in green code for amino acids as part of family boxes in the modern genetic code; those named in grey are the complementary anticodons for family-box triplets (with identical total GC compositions). Family box codons are thought to have been employed to recruit the first amino acids into a nascent coded peptide synthesis system, and the TPR RNA products are 3.8-fold enriched in them. There are contributions from both nucleobase composition (random sequences with identical nucleobase composition give a milder 2.8-fold preference for family box codons sequence) and triplet sequence preferences (there is just a 1.6-fold preference for family box anticodons). As may be expected based upon their composition, triplets corresponding to stop codons were rarely incorporated.

## Methods

### RNA preparation

**Method 1:** The indicated DNA oligonucleotide and oligonucleotide ‘HDVrt’ were mutually extended upon one another using three thermocycles with GoTaq Hot-start DNA polymerase (Promega). This generated a double-stranded DNA comprising the DNA sequence of the desired RNA oligonucleotide, downstream of a T7 promoter sequence, but upstream of the DNA sequence of the HDV ribozyme. This DNA was QIAQuick column purified (Qiagen) and transcribed in the presence of guanosine monophosphate (15 ng/µl DNA, 20 mM MgCl_2_, 50 mM Tris·HCl pH 7.9, 10 mM DTT, 2 mM spermidine, 6.25 mM of each of the four NTPs, 0.01 U/µl inorganic pyrophosphatase (Thermo Scientific), 20 ng/µl T7 RNA polymerase). During overnight transcription at 37°C, co-transcribed HDV ribozyme cleaved itself off from the RNA 3’ end to yield uniform 3’ termini; adding urea to 6 M and incubating for a further hour at 37°C enhanced this cleavage^42^. After PAGE purification the RNA was treated with polynucleotide kinase (NEB) to remove the residual 2’, 3’-cyclic phosphate before acid phenol/chloroform extraction to remove enzyme and ethanol precipitation to yield the final oligonucleotide.

**Method 2:** Ribozymes were prepared as per method 1 above, but the DNA transcription template (with the ribozyme-encoding sequence between a T7 promoter and HDV ribozyme) was generated by PCR. Transcription included excess guanosine monophosphate (GMP, 10 mM, with only 2 mM of each of the four NTPs) to yield 5’ monophosphorylated RNAs that avoid participation as substrates in ribozyme-catalysed ligation.

**Method 3:** Triphosphorylated triplets and oligomers were prepared as in ref. 4. Briefly, DNA oligonucleotide ‘5T7’ and the relevant templating DNA were mixed in transcription buffer at 1.5 µM to give a double-stranded T7 promoter with a downstream 5’ overhang sequence encoding the desired oligonucleotide (see ref. 4 for details of overhang choice for each triplet). The only nucleotides added to the transcription buffer were 2.4 mM of NTP for each nucleotide present in the desired product. The correct triplet product was then purified from a 30% polyacrylamide 3M urea gel and, after precipitation in 85% ethanol, its concentration was determined by UV absorbance and extinction coefficients calculated using OligoCalc^40^. ^ppp^NNN comprises an equimolar mix of the 64 possible triplet sequences.

**Method 4:** RNA adapter for RNA sequencing was transcribed as in method 1, but from DNA generated by mutual extension with oligonucleotide ‘5T7’ instead of ‘HDVrt’. As a result there was no HDV cleavage step in transcription. After PAGE purification, the RNA was treated with alkaline phosphatase (rSAP, NEB) instead of PNK to remove 5’ phosphorylation and prevent adapter concatenation during adapter ligation.

**Method 5:** RNA library was prepared as in method 4, but included GMP in transcription as in method 2, and omitted any alkaline phosphatase treatment after PAGE purification.

**Method 6: RNA duplex preparation.** All RNA duplexes were prepared by native PAGE purification. 600 pmol of each complementary RNA were annealed in 60µl of 0.1 M NaCl, 5 mM tris·HCl pH 7.4 (80°C 1 s, 0.1°C/s to 4°C, kept on ice until loading with glycerol added to 12%). After 0.5× TBE 20% PAGE separation (run at 10 W for 1 h) and excision of the duplex band (identified by comparison to single-strand controls), duplex was eluted into 0.1 M NaCl, 2 mM tris·HCl pH 7.4 overnight at 4°C. After removal of gel fragments by passage through a Spin-X 0.22 µm cellulose acetate filter (Costar), salts eluted from the gel were removed by applying the eluate to a 3 kDa molecular weight cut-off filter (Amicon Ultra-0.5) and filtering three times, diluting the concentrate with 0.1 M NaCl, 2 mM tris·HCl before each time. The final concentrate was supplemented with 0.1% Tween-20 and stored in a DNA LoBind microfuge tube (Eppendorf) in the fridge, where it showed no evidence of separation after > 6 months storage. Strand separation of duplexes was measured in a fluorescence/quench assay as described in Fig. S2.

### Ribozyme RNA polymerase assay

Standard (non-replicative) primer extension assays were conducted in 5 µl extension buffer (final concentrations 0.1 M MgCl_2_, 50 mM tris·HCl pH 8.3, 0.05% Tween-20, 100 nM of each primer, 2.5 µM of each triplet/oligomer, 0.5 µM ribozyme). To set up, ribozyme was annealed (80°C 2 min, 17°C 10 min) in 1 µl water. Meanwhile, 0.5 pmol of template or duplex was incubated in 2 µl 0.05% Tween-20 (+ 5 mM HCl for ‘acidified’ reactions) at 25°C for 10 min. All other reaction components (in 2 µl) were mixed with ribozyme then template fractions, before flash-freezing in liquid N_2_ (20 s) and incubation at −7°C in an LTC4 refrigerated bath (Grant Instruments).

To measure the effect of delayed triplet and/or TPR addition, duplex or template was acidified as above, then a neutralisation mix was added with or without the relevant triplets and primers. This neutralisation mix also contained enough MgCl_2_, tris·HCl pH 8.3, and Tween-20 to yield their final concentrations above. The resulting volume (2.5-3.7 µl) was immediately flash-frozen in liquid N_2_ and incubated at −7°C to allow the eutectic phase to thaw out, during which time the template strand(s) reannealed or became coated with triplets.

After the indicated delay, more buffer (with identical MgCl_2_/tris·HCl pH 8.3/Tween-20 concentrations) containing the TPR (with any missing triplets and primers) was added on top of the frozen volume to a final volume of 5 µl at −7°C. Overall, this now had an identical composition to the standard reactions, but the buffer with TPR froze as a distinct layer on top of the original ice; the eutectic phases of the layers, however, become contiguous, allowing TPR (+/- triplets) to diffuse into the lower layer containing the template (as previously reported^22^). Reactions were then incubated at −7°C for 48 h to allow primer extension; the fraction of primers extended was then used to deduce available template levels (and thus the degree of strand reannealing) as follows.

Data points with extension >0.1% were used to deduce reannealing rates in the eutectic phase. Even in standard reactions, primer extension is not complete, and (independent of strand reannealing) a 2-layer reaction with triplets separated from template gave three-fold less primer extension than when triplets began in the same layer as the template (Fig. 1c). Therefore, primer extension efficiencies on duplex were first divided by the average efficiency with template alone (from equivalent reactions with or without triplets at neutralisation - or a geometric mean thereof when half were present). Efficiencies were then converted into free strand concentrations by estimating the eutectic phase volume (see below). As triplets/TPR do not immediately complete diffusion between reaction layers, there is a hidden lag phase and extrapolated initial efficiencies would vary between different conditions. Nonetheless subsequent reannealing rates could be estimated using the changes observed between delays of different lengths.

### Replication cycling

Iterative replication of RNA was undertaken in 0.5 ml microfuge tubes containing 125 µl of replication buffer (standard composition: 0.4 mM MgCl_2_, 1.8 mM KCl, 1 mM CHES·KOH pH 9.0, 0.01% Tween-20, 20 nM TPR, 100 nM of each triphosphorylated triplet, 100 nM each of any RNA primers/oligomers). Up to 1 pmol of starting RNA template or duplex to be amplified was added per reaction (see Fig. S6 for discussion of this parameter). Reactions were prepared at room temperature, and cycling was begun with an initial denaturing step: 0.75 µl of 0.1 M HCl was added (+0.6 mM in the reaction, overwhelming the CHES buffer), the reaction was vortexed, and then incubated for 2 minutes on a thermocycler pre-heated to 80°C. Then 0.75 µl of 0.1 M KOH was added, the reaction briefly vortexed, and plunged in liquid N_2_ for 20 s to flash-freeze, before incubation at −7°C for ∼24 h. Another cycle was initiated by thawing the reactions at room temperature before HCl addition etc.

The fold-concentration of solutes within the eutectic phase upon freezing of extension buffer or replication buffer was calculated by estimating eutectic phase volume as a fraction of the ice, as described previously^22^. Briefly, a sample was prepared with the ionic composition of the target reaction (for extension buffer) or a 50-fold concentrated version (for replication buffer), alongside a gradually more concentrated set of samples with the same amount of buffer components but lower reaction volumes. These were then flash-frozen and incubated at −7°C to allow eutectic phase equilibration. If the volume of a concentrated sample was less than that of the eutectic phase of the parent sample, no ice phase would be present at equilibrium. This transition occurred in a 10-fold concentrated sample for extension buffer, and an 8.8-fold concentrated sample for 50× replication buffer. Assuming eutectic phase composition is the same independent of starting volume (i.e. uniform freezing point depression from solutes), linear concentration factors were applied (i.e. 440-fold concentration from freezing replication buffer at −7°C).

Open-ended replication cycling involving serial dilution proceeded as shown in the relevant figures (Fig. 2b, Fig. 3a, Fig. 5a). The cycling protocol was designed to create an appropriate replication burden (2-3 fold amplification in 4-5 cycles), whilst maintaining the KCl concentration (increasing from pH cycling) within the 1.8-4.2 mM range. To dilute, an acidified sample was neutralized post-heating by the combined addition of chilled KOH and fresh reaction mix, then flash frozen and incubated at −7°C. After this incubation, the reaction was thawed and a 125 µl aliquot transferred to a fresh tube for further cycling. The remnant sample was retained for analysis.

In some lanes of Fig. 4d, half of the TPR had been degraded beforehand by incubation at 80°C for 8 min in 20 mM MgCl_2_/30 mM KCl/50 mM CHES·KOH pH 9.0/0.02% Tween-20; the remaining reaction components were then added to this mixture to restore the standard reaction composition before cycling began.

### Denaturing gel electrophoresis

For gel analysis of RNA synthesis in extension buffer, samples were quenched with excess EDTA over Mg^2+^, and urea to 6M. Where a specific template sequence was included in the reaction, a 10-fold excess of a complementary RNA was also provided to outcompete product:template rehybridization. Samples were denatured (94°C, 5 min), cooled and separated on 20% acrylamide/8 M urea/TBE gels. For analysis of RNA synthesis in replication buffer, 125 µl reaction aliquots were first ethanol precipitated (85% ethanol final concentration, with glycogen carrier) before resuspension in water and addition of EDTA and urea as above.

Reactions in Fig. S8b with biotinylated primer were stopped with excess EDTA and mixed with 2 volumes of BWBT buffer (200 m NaCl, 10 mM tris·HCl pH 7.4, 1 mM EDTA, 0.1% Tween-20), then biotinylated primers and products were bound to MyOne C1 streptavidin-coated microbeads (Invitrogen) prewashed thrice in BWBT. These were then washed with BWBT, templates were removed with a 50 mM NaOH/1 mM EDTA/0.1% Tween-20 wash, beads were neutralized with a BWBT + 100 mM tris·HCl pH 7.4 wash, before a final BWBT wash and resuspension in 95% formamide/25 mM EDTA. After heating at 94°C for 5 min to denature and detach primers from beads, the supernatant was subjected to denaturing PAGE to separate the primers.

Gels were scanned on a Typhoon FLA-9000 imager (GE) at different wavelengths for each fluorophore-labelled primer. The gel of primerless extensions in Fig. 5b was pre-stained with SYBR Gold (Invitrogen) before scanning. Gel bands were quantified using ImageQuant analysis software; backgrounds were drawn between peak troughs or using adjacent negative controls. Band intensities of the total extension products and unextended primers were used to calculate product distribution and hence extension yields of e.g. ‘full-length’ (full-length band only), ‘reaching full-length’ (including longer bands), or ‘primer extension’ (at least 1-3 triplet addition) products as indicated in figures.

Oligonucleotides were purified from PAGE gels by excision of bands identified by UV shadowing or alignment with a fluorescence scan. Bands were crushed and oligonucleotides therein eluted into 10 mM tris·HCl pH 7.4. Gel fragments were removed from the supernatant by passage through a Spin-X 0.22 µm cellulose acetate filter (Costar) and the supernatant precipitated in 73% ethanol (ribozymes or long oligonucleotides) or 85% ethanol (oligonucleotides < 8 nt).

### Sequencing of primerless fragment syntheses

To understand the synthesis pathways of ribozyme fragments (Fig. S5), polymerisation products made on templates without primers by the t5^+1^ TPR^4^ were eluted and sequenced. Reaction profiling was carried out on ribozyme reactions set up with 4 pmol of biotinylated template (Bbio, ybio or dbio encoding the β^+^, γ^+^ or δ^+^ fragments^4^ of t5 respectively) prebound to 40 µg MyOne C1 microbeads, 2 pmol of t5^+1^, and 50 pmol of each target fragment’s constitutive triplet in 10 µl of 2× extension buffer, supplemented with 0.15% Tween-20. Reactions were frozen at −7°C and stopped after 4 (Bbio, ybio) or 11 (dbio) days by vortexing until thawed, whereafter all steps were carried out on ice to maximise the retention of any short products hybridised to templates.

Beads were twice washed with ice cold BWBTMg (200 mM NaCl, 10 mM tris·HCl pH 7.4, 20 mM MgCl_2_, 0.1% Tween-20) to remove substrates and ribozyme whilst maintaining duplex annealing, changing to a new pre-chilled tube in between. Beads were then resuspended in 5 µl 10 mM EDTA, 0.1% Tween-20, before addition of 5 µl 80 mM NaOH to elute products from templates. This supernatant was removed after 1 min and neutralised with a mix of 0.4 µl 1 M tris·HCl pH 7.4 and 0.4 µl 1 M HCl.

Sequence analysis began with 3’ adaptor ligation: 2.7 µl of the sample of eluted extension products were included in a 10 µl T4 RNA ligase 2 truncated KQ reaction (1× T4 RNA ligase buffer (NEB), 2 µM IllLigBio adapter pre-adenylated using a 5’ adenylation kit (NEB), 20-200 nM ^ppp^UGAAUG hexamer standard, 15% PEG, 20 U/µl of T4 RNA ligase 2 truncated KQ (NEB), 10°C overnight). RNA-adapter ligation products were purified by denaturing PAGE, then eluted and ethanol precipitated with a glycogen carrier as described above. After resuspension in 5 µl water, 5’ triphosphates on the synthesis products were converted into 5’ monophosphates: samples were treated with recombinant shrimp alkaline phosphatase (NEB) (10 µl reaction in 1× NEB buffer 2.1, 37°C 30 min, 65°C 5 min), these reactions were made up to 15 µl including 0.5 µl 10× NEB 2.1, 1.5 µl 10 mM ATP, 0.75 µl 0.1 M DTT and 0.3 U/µl T4 Polynucleotide kinase (NEB), and then incubated at 37°C 30 min, 65°C 20 min.

After addition of a monophosphate, 6 µl aliquots were 5’ adapter ligated (25 µl of 1× T4 RNA ligase buffer (NEB), 1 mM ATP, 0.4 µM p10Illligup adaptor, 25% PEG, 0.5 U/µl T4 RNA ligase 1, 25°C 2 hr, 16°C overnight). Adapters contained random sequence stretches and a short mutually-complementary region to maximise ligation efficiency and generality^39^. 2.5 µl aliquots were subsequently reverse transcribed and PCR amplified in a 50 µl SuperScript III one-step RT-PCR System (Invitrogen), using 0.4 µM barcoded P5XGGGP10 and P3Illrt primers. PCR products were agarose gel purified using a Monarch gel extraction kit (NEB) and sequenced on an Illumina HiSeq.

Using the Galaxy web platform at the public server usegalaxy.org^43^, the first three positions (of random sequence) and any positions beyond 120 nt were trimmed off the sequencing reads, before quality filtering (requiring >90% of positions with >Q=25) and conversion to .fasta format. Sequences possessing AGATCGGAAGAG from the 3’ adapter were then retained. This motif, the preceding 13 positions and all downstream positions were then trimmed off to remove the 3’ adapter sequence before the data was split by the identity of the first six ‘barcode’ positions of each sequence, also requiring an intact 5’ adapter sequence: XXXXXXGGGCTGCCAACCGACCGACTATC. The first 42 positions of each sequence were then trimmed to fully remove the 5’ adapter, yielding raw data files of insert sequences (‘Btri4_GRC.fasta’, ‘ytri4_GRC.fasta’, ‘dtri11_GRC.fasta’), with 130,000 – 780,000 reads per test sample. In parallel negative control reactions were set up without triplets, and processed and analysed similarly (‘Bneg_GRC.fasta’, ‘yneg_GRC.fasta’, ‘dneg_GRC.fasta’).

Using custom script ‘intermaster1.2’ (Table S2), reads were then mapped against a list of all possible species for their respective template generated using custom script ‘specify’ (Table S2), allowing for one mismatch in the three 5’ and three 3’ positions, or one mismatch anywhere for hexamer reads. This produced counts of each oligonucleotide product (‘counts_compiled_2.2.xlsx’) which were then normalised by the count of the spiked-in ^ppp^UGAAUG hexamer, reduced by reads from negative control samples (comprising 0.03 - 16% of the number of sample reads), and converted to fmol or amol values of each product made per pmol of template, assuming a linear scaling of reads with concentration. These data were used to generate the fragment synthesis pathways as described in Fig. S5b.

### Sequencing of γ fragment replication products

γ^+^ synthesis products were purified by PAGE as described in Fig. S8a & b. After 85% ethanol precipitation with glycogen carrier, these products were ligated to a 3’ adapter (1× T4 RNA ligase buffer (NEB), 10% PEG-8000, 10 U/µl T4 RNA Ligase 2 truncated KQ (NEB), 2 µM HDVLig (pre-adenylated using a 5’ adenylation kit (NEB)), 10°C overnight, 65°C 10 minutes). This ligation was spiked at 2% into a SuperScript III/Platinum Taq rtPCR reaction (Invitrogen) with HPγseq and HDVrt primers. To add sequencing tags, this reaction was spiked at 1% into a GoTaq HotStart (Promega) PCR reaction with primers P3HDV and P5XHPγn, and the desired product was agarose gel purified using a Monarch gel extraction kit (NEB) and sequenced on an Illumina MiSeq.

The Galaxy web platform was used for initial processing of reads as described above, with sequences possessing GGGTCGGCATGGCATC from the 3’ adapter retained, before this motif and all downstream positions were trimmed off. The data was split by the sequence: XXXXXXGACTCTTCGGAGTCTTCGGAT containing a barcode region and the FITCγSeq primer sequence, yielding raw data files of extended primers (‘1minus.fasta’, ‘1yD.fasta’, ‘5minus.fasta’, ‘5yd.fasta’), with 59,000 – 74,000 reads per test sample. The first 27 positions of each sequence were then trimmed to fully remove the 5’ primer sequence, and any extension products shorter than full-length (24 nt) were discarded, yielding full-length extension product data files (‘1minusFL+.fasta’, ‘1yDFL+.fasta’, ‘5minusFL+.fasta’, ‘5ydFL+.fasta’). The degree of identity of 24-nt sequences with the target sequence GCAGAGGCGGCAGCCTTCGGTGGC in each sample was counted by custom script ‘tally’ (Table S2), and these distributions were compared to the identity distribution of sequences generated from a random mix of the substrate triplets present (built using sequences generated by custom script ‘24ntrandom’ (Table S2)), as detailed in source data file ‘Error tallies weighted.xlsx’ and shown in Fig. S8c.

### Sequencing of triphosphorylated amplification products

To identify TPR-synthesised RNA products from cycling in the absence of primers with ^ppp^NNN, 3’ adapters were ligated to RNAs purified from the gel in Fig. 5b (1× T4 RNA ligase buffer (NEB), 1.6 µM IllLigBio adapter pre-adenylated using a 5’ adenylation kit (NEB), 15% PEG, 16 U/µl of T4 RNA ligase 2 truncated KQ (NEB), 10°C 20 h, 65°C 12 min). The reaction was then supplemented with 40 mM tris·HCl pH 8.3 and 0.5 U/µl RppH (NEB) before incubation (37°C 80 min) to convert 5’ triphosphates to 5’ monophosphates before purification of ligated products by denaturing PAGE. These were eluted and precipitated before ligation to 5’ adapters (1× T4 RNA ligase buffer (NEB), 0.5 µM p10IllLigup adapter, 25% PEG-8000, 1 mM ATP, 2 U/µl T4 RNA ligase 1, 25°C 2 hr, 16°C 9 h, 65°C 15 min). This ligation was spiked at 10% into a SuperScript III/Platinum Taq rtPCR reaction (Invitrogen) with 0.4 µM P3Illrt and P5XGGGP10 primers, and the desired products were agarose gel purified using a Monarch gel extraction kit (NEB) and sequenced on an Illumina HiSeq.

Sequences were processed using the FASTX Toolkit^44^ version 0.0.13. The first three positions (of random sequence) were trimmed off the sequencing reads, before quality filtering (requiring >95% of positions with >Q=30) and conversion to .fasta format. Sequences possessing AGATCGGAAGAG from the 3’ adapter were then retained. This motif, the preceding 13 positions and all downstream positions were then trimmed off to remove the 3’ adapter sequences. The data was split by the identity of the first six ‘barcode’ positions of each sequence attached to an intact 5’ adapter sequence: XXXXXXGGGCTGCCAACCGACCGACTATC. The first 42 positions of each sequence were then trimmed to fully remove the 5’ adapter, yielding raw data files of insert sequences (’RZ5N.fasta’, ’N17L.fasta’, ’N73L.fasta’, ’N73H.fasta’, ’F201N.fasta’, ’F205N.fasta’, ’FN17L.fasta’, ’FN73L.fasta’, and ’FN73H.fasta’, corresponding to samples A-I respectively), with 39,000-640,000 reads per sample.

Sequence_lengths.py followed by Plot_sequence_lengths.R were used to plot sequenced product length distributions (Fig. S9b). Then custom code in Alignments.ipynb was used to exclude non-triplet register products (Fig. S9c) and products > 3 nt outside the size range excised from the gel as in Fig. S9a. To classify product sequences by alignment to ribozyme (+) and (−) sequences (Fig. S9d), the Biopython local alignment tool was used (pairwise2.align.localms) as part of custom code Alignments.ipynb. The gap penalty was set to −1000, and a score of 1 was given to each matching nucleotide and 0 for every mismatch. Dividing the score by the length of the given sequence yielded the fractional identity with the ribozyme, and site and degree of maximal fractional identity was recorded. Sequences above a fractional identity threshold (Fig. S9c) were classified as ‘aligned’. This script was adapted to only align the 4^th^-12^th^ nt of each product to perform length-independent classification in Fig. S9e (Alignments.ipynb & Plot_alignment_count.R).

To generate the map of alignment sites in Fig. 5h (Plot_alignment_ribozyme.R), a score was given to positions in the ribozyme sequence for each ‘aligned’ product. The score was corrected for the yield of different reactions, taking into account the intensity of product bands on the gel and the total number of sequences that went into the alignment in each pool. Furthermore, if a product aligned to more than one of the four possible ribozyme sequences (5TU, t1 and complements), the score increase was shared between the relevant sites to avoid inducing bias at regions of high sequence similarity between 5TU and t1.

Custom scripts (Alignments.ipynb / Family_box_codons.py / Length_vs_identity.py) were used to generate randomised pools of sequences for control analyses in Fig. S9g / Fig. 5d / Fig. 5e & Fig. S9f and plotted using Plot_alignment_count.R / Plot_family_box_codons.R / Plot_length_vs_identity.R, respectively. For Fig. 5e, S9f and S9g, the sequences were randomised having the same triplet composition and length distribution as the comparison pools, while for Fig. 5d, only the length distribution was maintained and otherwise completely random sequences were chosen. Custom script (Family_box_codons.py and Plot_family_box_codons.R) was used to count contiguous family box triplets in product sequences for Fig. 5d. Length_vs_identity.py allows counting the maximum identity of a product sequence upon alignment with the ribozyme, and comparison to its total length, which is plotted with Plot_length_vs_identity.R (Fig. 5e and S9f).

The uniqueness of product sequences was assessed with custom script Uniqueness.py and plotted with Plot_uniqueness.R (Fig. S10). Complementarity of sequences was analysed with Complementarity.py and plotted with Plot_complementarity.R (Fig. S10). The nucleotide composition of triplets was plotted with Plot_nucleotide_triplets.R with files coming from Alignments.ipynb (Fig. 5g).

## Acknowledgements

This work was supported by the Medical Research Council (MRC) as part of United Kingdom Research and Innovation (also known as UK Research and Innovation (UKRI)) (MRC program grant MC_U105178804) (JA, TA, JFC, SK, EG, PH), a Royal Society University Research Fellowship (URF\R1\201271) (JA) and a grant from the Volkswagen Foundation (96 755) (EG). The authors thank E.L. Kristoffersen, B.T. Porebski, and J.D. Sutherland for helpful discussions. For the purpose of open access, the MRC Laboratory of Molecular Biology has applied a CC BY public copyright license to any Author Accepted Manuscript version arising.

## Contributions

Project was conceived and designed by JA & PH. JA designed, performed and analysed experiments, together with SK (design of template A and replication buffer), JFC (design, execution and analysis of synthesis pathways (Fig. S5)), EG (TPR construct development and sequence processing), and TA (analysis and presentation of triphosphorylated amplification reactions (Fig. 5, Fig. S9, Fig. S19, Supplementary Table 3)). JA and PH wrote the paper and all authors commented on the manuscript.

## References

1. Szostak, J.W. The Eightfold Path to non-enzymatic RNA replication. J. Syst. Chem. 3 (2012).

2. Zhang, S.J., Duzdevich, D., Ding, D. & Szostak, J.W. Freeze-thaw cycles enable a prebiotically plausible and continuous pathway from nucleotide activation to nonenzymatic RNA copying. Proc. Natl Acad. Sci. USA 119, e2116429119 (2022).

3. Leveau, G., et al. Enzyme-Free Copying of 12 Bases of RNA with Dinucleotides. Angew. Chem. Int. Ed. Engl. 61, e202203067 (2022).

4. Attwater, J., Raguram, A., Morgunov, A.S., Gianni, E. & Holliger, P. Ribozyme-catalysed RNA synthesis using triplet building blocks. eLife 7, e35255 (2018).

5. Portillo, X., Huang, Y.T., Breaker, R.R., Horning, D.P. & Joyce, G.F. Witnessing the structural evolution of an RNA enzyme. eLife 10, e71557 (2021).

6. Cojocaru, R. & Unrau, P.J. Processive RNA polymerization and promoter recognition in an RNA World. Science 371, 1225–1232 (2021).

7. Tjhung, K.F., Sczepanski, J.T., Murtfeldt, E.R. & Joyce, G.F. RNA-Catalyzed Cross-Chiral Polymerization of RNA. J. Am. Chem. Soc. 142, 15331–15339 (2020).

8. Freier, S.M. et al. Improved free-energy parameters for predictions of RNA duplex stability. Proc. Natl. Acad. Sci. USA 83, 9373–7 (1986).

9. Li, Y. & Breaker, R.R. Kinetics of RNA Degradation by Specific Base Catalysis of Transesterification Involving the 2’-Hydroxyl Group. J. Am. Chem. Soc. 121, 5364–5372 (1999).

10. Mariani, A., Bonfio, C., Johnson, C.M. & Sutherland, J.D. pH-Driven RNA Strand Separation under Prebiotically Plausible Conditions. Biochemistry 57, 6382–6386 (2018).

11. Ianeselli, A. et al. Water cycles in a Hadean CO2 atmosphere drive the evolution of long DNA. Nature Physics 18, 579–585 (2022).

12. Ianeselli, A., Mast, C.B. & Braun, D. Periodic Melting of Oligonucleotides by Oscillating Salt Concentrations Triggered by Microscale Water Cycles Inside Heated Rock Pores. Angew. Chem. Int. Ed. Engl. 58, 13155–13160 (2019).

13. Ianeselli, A. et al. Physical non-equilibria for prebiotic nucleic acid chemistry. Nature Reviews Physics (2023).

14. Salditt, A. et al. Ribozyme-mediated RNA synthesis and replication in a model Hadean microenvironment. Nat. Commun. 14, 1495 (2023).

15. He, C., Lozoya-Colinas, A., Gallego, I., Grover, M.A. & Hud, N.V. Solvent viscosity facilitates replication and ribozyme catalysis from an RNA duplex in a model prebiotic process. Nucleic Acids Res. 47, 6569–6577 (2019).

16. Lozoya-Colinas, A., Clifton, B.E., Grover, M.A. & Hud, N.V. Urea and Acetamide Rich Solutions Circumvent the Strand Inhibition Problem to Allow Multiple Rounds of DNA and RNA Copying. Chembiochem 23, e202100495 (2022).

17. Zhou, L. et al. Non-enzymatic primer extension with strand displacement. eLife 8, e51888 (2019).

18. Kristoffersen, E.L., Burman, M., Noy, A. & Holliger, P. Rolling circle RNA synthesis catalyzed by RNA. eLife 11, e75186 (2022).

19. Horning, D.P. & Joyce, G.F. Amplification of RNA by an RNA polymerase ribozyme. Proc. Natl. Acad. Sci. USA 113, 9786–91 (2016).

20. Bare, G.A.L. & Joyce, G.F. Cross-Chiral, RNA-Catalyzed Exponential Amplification of RNA. J. Am. Chem. Soc. 143, 19160–19166 (2021).

21. McRae, E.K.S., et al. Cryo-EM structure and functional landscape of an RNA polymerase ribozyme. bioRxiv, 2022.08.23.504927 (2022).

22. Attwater, J., Wochner, A., Pinheiro, V.B., Coulson, A. & Holliger, P. Ice as a protocellular medium for RNA replication. Nat. Commun. 1, 76 (2010).

23. Markham, N.R. & Zuker, M. DINAMelt web server for nucleic acid melting prediction. Nucleic Acids Res. 33, W577–81 (2005).

24. Attwater, J., Wochner, A. & Holliger, P. In-ice evolution of RNA polymerase ribozyme activity. Nat. Chem. 5, 1011–8 (2013).

25. Kreysing, M., Keil, L., Lanzmich, S. & Braun, D. Heat flux across an open pore enables the continuous replication and selection of oligonucleotides towards increasing length. Nat. Chem. 7, 203–8 (2015).

26. Chargaff, E. Some recent studies on the composition and structure of nucleic acids. J. Cell. Physiol. 38, 41–59 (1951).

27. Rosenberger, J.H. et al. Self-Assembly of Informational Polymers by Templated Ligation. Physical Review X 11, 031055 (2021).

28. Mulkidjanian, A.Y., Bychkov, A.Y., Dibrova, D.V., Galperin, M.Y. & Koonin, E.V. Origin of first cells at terrestrial, anoxic geothermal fields. Proc. Natl. Acad. Sci. USA 109, E821–30 (2012).

29. Cousins, C.R. et al. Biogeochemical probing of microbial communities in a basalt-hosted hot spring at Kverkfjoll volcano, Iceland. Geobiology 16, 507–521 (2018).

30. Keil, L.M.R., Moller, F.M., Kiess, M., Kudella, P.W. & Mast, C.B. Proton gradients and pH oscillations emerge from heat flow at the microscale. Nat. Commun. 8, 1897 (2017).

31. Morasch, M. et al. Heated gas bubbles enrich, crystallize, dry, phosphorylate and encapsulate prebiotic molecules. Nat. Chem. 11, 779–788 (2019).

32. Mutschler, H., Wochner, A. & Holliger, P. Freeze-thaw cycles as drivers of complex ribozyme assembly. Nat. Chem. 7, 502–8 (2015).

33. Roberts, S.J., Liu, Z. & Sutherland, J.D. Potentially Prebiotic Synthesis of Aminoacyl-RNA via a Bridging Phosphoramidate-Ester Intermediate. J. Am. Chem. Soc. 144, 4254–4259 (2022).

34. Zhou, L., Ding, D. & Szostak, J.W. The virtual circular genome model for primordial RNA replication. RNA 27, 1–11 (2021).

35. Jain, N. et al. Transcription polymerase-catalyzed emergence of novel RNA replicons. Science 368, 153–163 (2020).

36. Grosjean, H. & Westhof, E. An integrated, structure- and energy-based view of the genetic code. Nucleic Acids Res. 44, 8020–40 (2016).

37. Borsenberger, V. et al. Exploratory studies to investigate a linked prebiotic origin of RNA and coded peptides. Chem. Biodivers. 1, 203–46 (2004).

38. Patel, B.H., Percivalle, C., Ritson, D.J., Duffy, C.D. & Sutherland, J.D. Common origins of RNA, protein and lipid precursors in a cyanosulfidic protometabolism. Nat. Chem. 7, 301–7 (2015).

## Supplementary References

39. Fuchs, R.T., Sun, Z., Zhuang, F. & Robb, G.B. Bias in Ligation-Based Small RNA Sequencing Library Construction Is Determined by Adaptor and RNA Structure. PLOS ONE 10, e0126049 (2015).

40. Kibbe, W.A. OligoCalc: an online oligonucleotide properties calculator. Nucleic Acids Res. 35, W43–6 (2007).

41. Cock, P.J. et al. Biopython: freely available Python tools for computational molecular biology and bioinformatics. Bioinformatics 25, 1422–3 (2009).

42. Rosenstein, S.P. & Been, M.D. Self-cleavage of hepatitis delta virus genomic strand RNA is enhanced under partially denaturing conditions. Biochemistry 29, 8011–8016 (1990).

43. Afgan, E. et al. The Galaxy platform for accessible, reproducible and collaborative biomedical analyses: 2016 update. Nucleic Acids Res. 44, W3–W10 (2016).

44. Hannon, G.J. FASTX-Toolkit. http://hannonlab.cshl.edu/fastx_toolkit (2010).

